# Generating 3D Models of Carbohydrates with GLYCAM-Web

**DOI:** 10.1101/2025.05.08.652828

**Authors:** Oliver C. Grant, Daniel Wentworth, Samuel G. Holmes, Rajan Kandel, David Sehnal, Xiaocong Wang, Yao Xiao, Preston Sheppard, Tobias Grelsson, Andrew Coulter, Grayson Miller, Bethany L. Foley, Robert J. Woods

**Affiliations:** Complex Carbohydrate Research Center and Department of Biochemistry and Molecular Biology, University of Georgia, 315 Riverbend Road, Athens, Georgia 30602, United States; National Centre for Biomolecular Research, Faculty of Science, Masaryk University, Brno, 625 00, Czech Republic; Hubei Key Laboratory of Agricultural Bioinformatics, College of Informatics, Huazhong Agricultural University, Wuhan, Hubei, China

## Abstract

The carbohydrate 3D structure-prediction tools (builders) at GLYCAM-Web (glycam.org) are widely used for generating experimentally-consistent 3D structures of oligosaccharides suitable for data interpretation, hypothesis generation, simple visualization, and subsequent molecular dynamics (MD) simulation. The graphical user interface (GUI) enables users to create carbohydrate sequences (e.g. DGalpb1-4DGlcpb1-OH) that are converted to 3D models of the carbohydrate structures in multiple formats, including PDB and OFF (AMBER software format). The resulting structures are energy minimized prior to download and online visualization. There are advanced options for selecting which shapes (rotamers) of the oligosaccharide to generate, and for creating explicitly solvated structures for subsequent MD simulation. The GLYCAM-Web builders integrate known conformational preferences of oligosaccharides, summarized here, and employ the GLYCAM forcefield for energy minimization with algorithms tailored for speed and scalability. Even for large oligosaccharides (100 residues, ~2100 atoms) a 3D structure is typically returned to the user in less than a minute.

---

Together with proteins, nucleic acids, and lipids, carbohydrates constitute a fundamental biomolecular building block. Comprised of monosaccharide residues and commonly referred to as oligosaccharides, polysaccharides or glycans, carbohydrates have unique characteristics that significantly impact their 3D structures, dynamics, and properties. First, monosaccharides can be connected in multiple ways; while two alanine residues form a single dipeptide, two glucopyranoses can form 10 chemically distinct disaccharides. Second, polysaccharides are often modified with chemical moieties such as sulfate, phosphate, or acetyl groups, which are essential for their biological function. For example, variations in the sulfation patterns of heparin regulate its anticoagulant activity^1^. Third, carbohydrates are not limited to the formation of linear polymers. The branching patterns create distinct 3D shapes that influence oligosaccharide shape and biological recognition^2,3^. The complexity of carbohydrate structures enables them to encode vast biological information. Lastly, although carbohydrates are decorated with hydroxyl groups, internal hydrogen bonding that might stabilize their 3D shapes is disrupted by hydrogen bonds with water^4^. As a result, glycans do not fold into well-defined secondary or tertiary structures, but instead adopt multiple conformations in solution, complicating their structural analysis^5^. Fortunately, the physical principles behind this behavior are well understood, and computational force fields that employ the pertinent physical properties for carbohydrates have been have been developed for each of the common biomolecular simulation packages, including GLYCAM/AMBER^6–8^, CHARMm^9–11^, and GROMOS^12–14^. Critically, glycans only adopt a small subset of the possible shapes for such a polymer. These shape preferences have been recognized since the early days of NMR spectroscopy^15,16^ and studied in detail computationally^5^. Thus, in contrast to other biopolymers, the strict structural preferences that determine glycan 3D shape have the benefit of enabling the 3D shapes of a glycan to be accurately predicted based solely on a knowledge of the glycan primary structure (i.e. the monosaccharide sequence, as well as the anomeric configuration and the linkage positions between the monosaccharides).

Despite the existence of established carbohydrate force fields, their practical implementation and adoption is hindered by the complexity of carbohydrate structure rules and specialized nomenclature to describe these non-linear polymers. These challenges can deter biophysicists and experimental scientists trained in protein study from examining carbohydrates, and correspondingly pose barriers for theoreticians. Developing carbohydrate-specific computational tools for use by those with limited domain-specific knowledge provides a mechanism to bridge the gaps between these fields and so accelerate adoption of sophisticated modeling methods for carbohydrates. For these reasons, GLYCAM-Web (www.glycam.org) was created in 2005 to provide a convenient online tool for predicting the 3D shapes of glycans, and has undergone continuous evolution to expand its capabilities, improve its breadth and accuracy, and respond to user feedback. Here we provide the first report of the current capabilities of the “carbohydrate-builders” available at GLYCAM-Web, as well as a discussion of the scientific logic that underpins carbohydrate 3D structure prediction.

## Results

### Types of builders on GLYCAM-Web

The various carbohydrate builders presented in **Table 1** provide a variety of ways to generate a carbohydrate sequence that are tailored to the experience level of the user. The point-and-click interface employs the Symbol Nomenclature for Glycans (SNFG)^17^ to help guide users familiar with this nomenclature (**Figure 1**), and was designed with the goal of minimizing the number of clicks necessary to build a sequence.

**Table 1.** The Carbohydrate Builder Interfaces Available at GLYCAM-Web.

| Builder Name | ShortLink | Description |
| --- | --- | --- |
| Point & Click<sup>a</sup> | glycam.org/cb | Generate a sequence by selecting residues and linkages. Right-to-left sequence entry. |
| Point & Click (ltr)<sup>a</sup> | glycam.org/cb/ltr | Generate a sequence by selecting residues and linkages. Left-to-right sequence entry. |
| 3D Structure Libraries | glycam.org/lib | Select sequences of common glycans. |
| Build-via-text | glycam.org/txt | Enter a sequence (type or cut-paste). |
| Build-via-url | glycam.org/url | Enter a sequence into the browser. |
| GAG Builder<sup>18</sup> | glycam.org/gag | Generate sequences of glycosaminoglycans. |

<sup>a</sup> Both left-to-right and right-to-left sequence entry are supported.

**Table 1.**
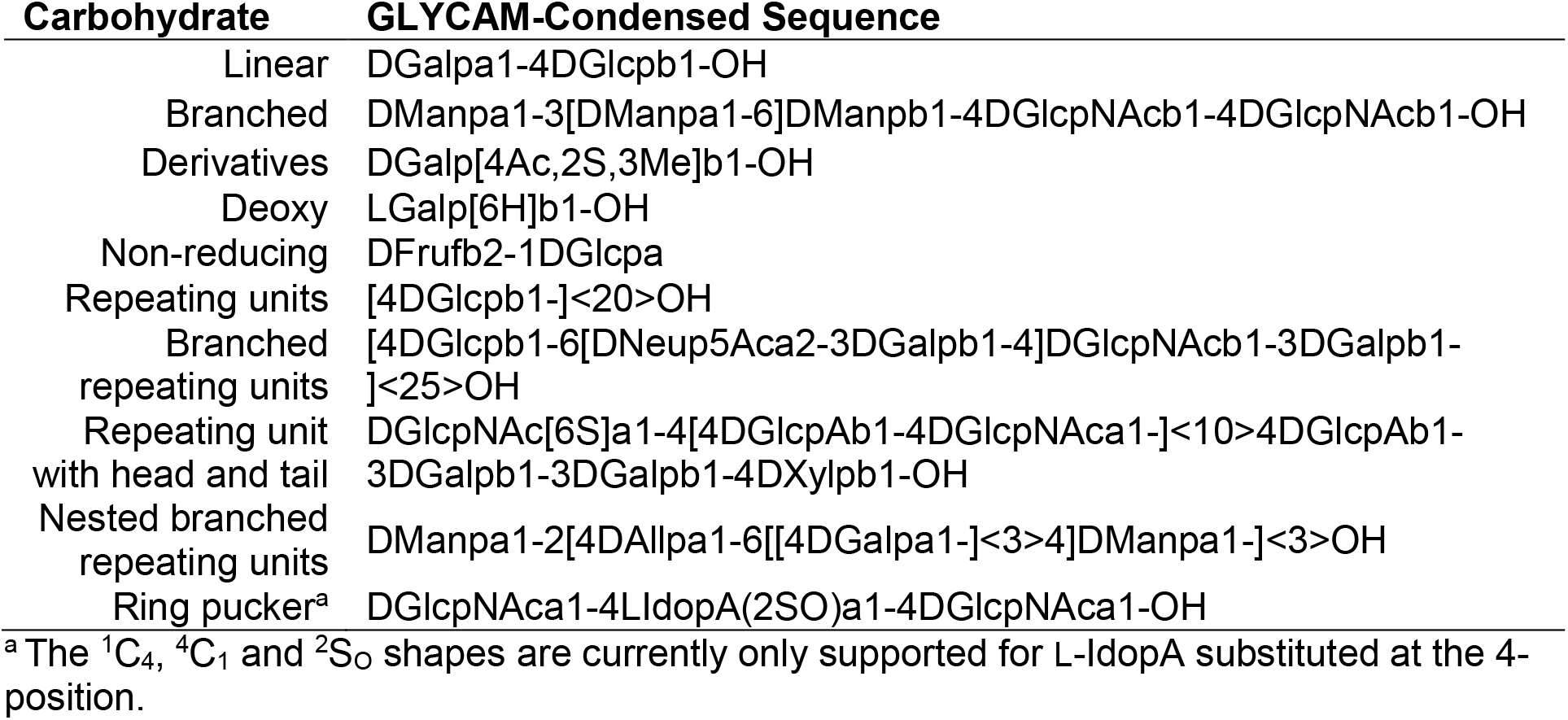
Examples of the available features in the GLYCAM condensed carbohydrate nomenclature.

**Figure 1.**
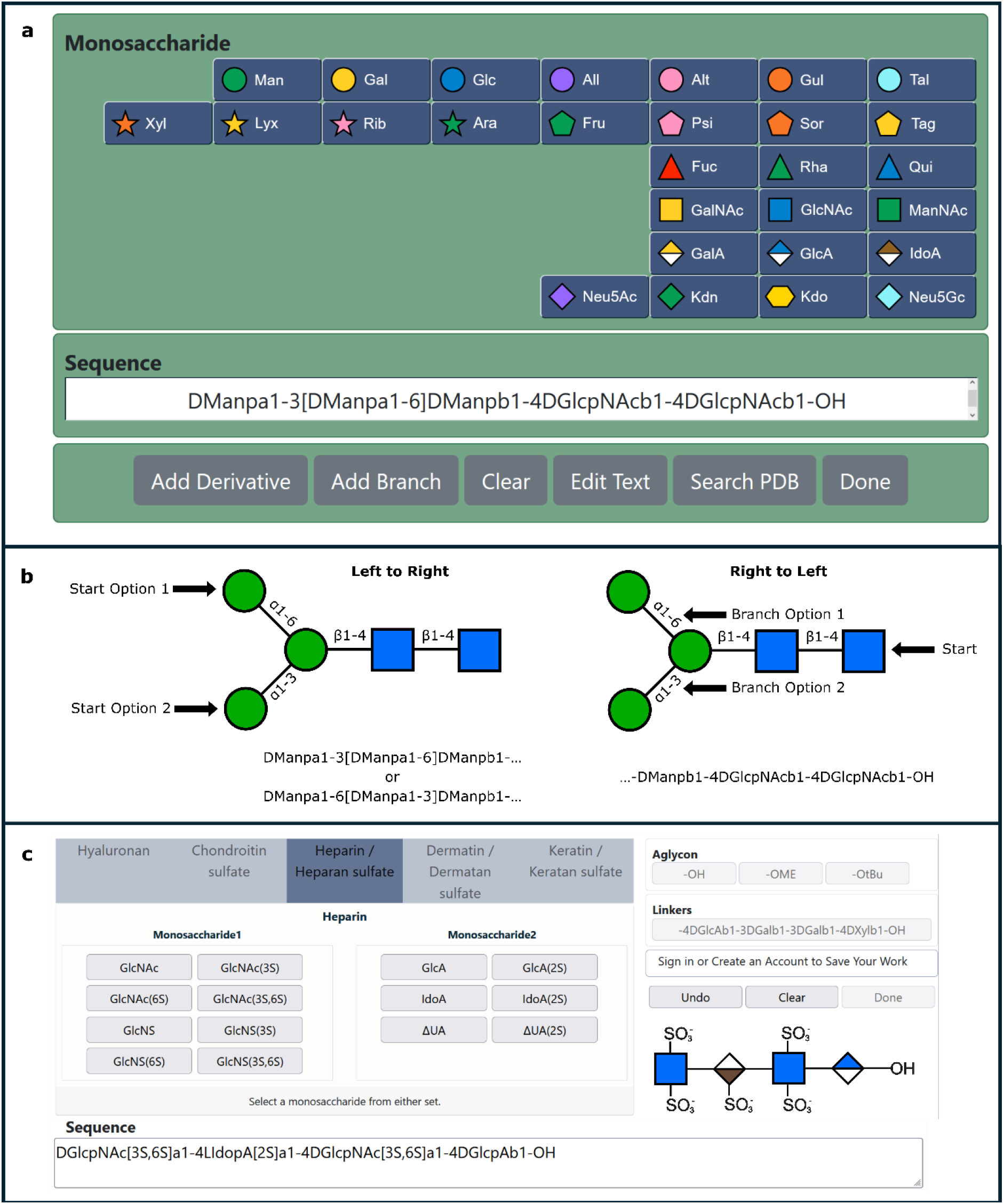
a) The point-and-click Carbohydrate Builder (CB) interface employs well-recognized SNFG^17^ symbols and permits sequence entry, addition of chemical derivatives, and creation of branches. It also permits searching of the protein data bank (PDB) for experimental data containing the specified carbohydrate. b) The oligosaccharide sequences can be entered from left to right (non-reducing to reducing end) or right to left (reducing to non-reducing end). c) The glycosaminoglycan builder^18^ integrates biological information to allow users to generate sequences for complex sulfated oligosaccharides with a minimum of clicks.

The 3D structure libraries at GLYCAM-Web contain common mammalian glycans, including high mannose *N-*glycans, as well as sialylated or fucosylated variants, and the entire set of 611 carbohydrates from the Consortium for Functional Glycomics glycan array (version 5.0). Users may select a structure from a library, and the tool automatically generates a 3D model. The build-via-text or build-via-url tools provide a convenient interface for more experienced users to insert and modify previously generated sequences by cutting-and-pasting a sequence, such as one selected from the 3D structure libraries. Build-via-text also supports manual editing of the GLYCAM condensed nomenclature.

Oligosaccharide sequences are normally specified in a left-to-right order with the reducing terminus at the rightmost position. However, some users new to the field reported that for branched oligosaccharides it could be counterintuitive to begin entering a sequence at an arbitrary non-reducing terminus and build towards the reducing terminus, thus both building directions are supported by GLYCAM-Web (**Figure 1**).

After requesting a sequence, users are either brought directly to a download page, or, if their sequence contains a flexible linkage with multiple potential rotamers, they are first taken to an options page to select which rotamers to generate (**Figure 2**). For visualization of the resulting 3D structures on the options and download pages, GLYCAM-Web implements the Mol* (“MolStar”) viewer^19,20^, which has integrated the “3D-SNFG” symbols^21^ for monosaccharides (**Figure 2**).

**Figure 2.**
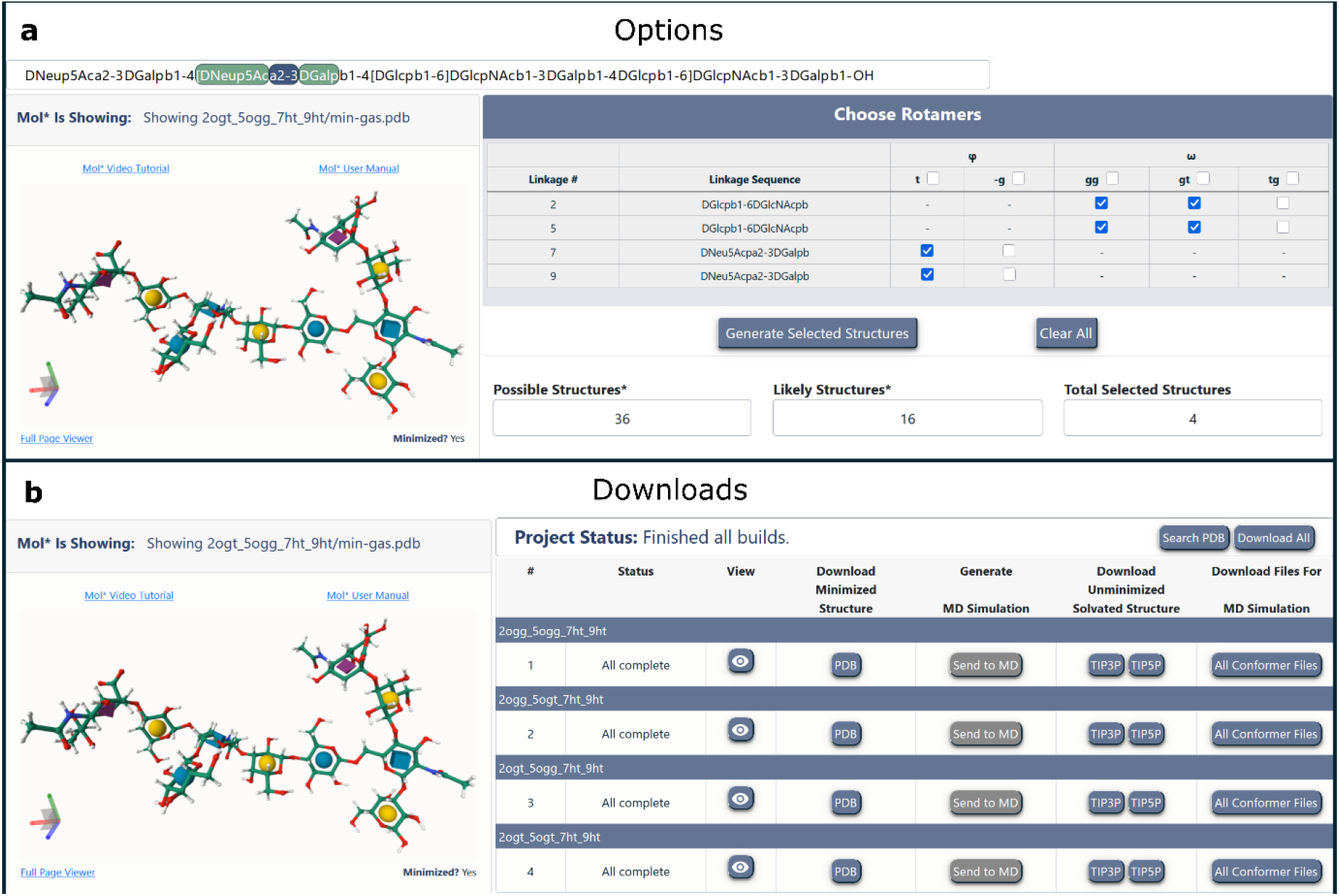
a) The options page allows the user to select which rotamers to generate for each glycosidic linkage that is known to populate more than one rotamer in solution. Hovering over a linkage in the options table highlights it in the sequence. Each combination of selected rotamers is generated. b) The download page displays the status of the energy minimization process for each resulting shape of the carbohydrate, and once minimization is complete the files become available for visualization and download individually or collectively in a zip file. The 3D shape of the carbohydrate is displayed in a side panel using the MolStar viewer^19,20^ with “3D-SNFG” symbols^21^ to aid in recognition of each monosaccharide.

### The GLYCAM-Web API

In addition to the user interfaces listed in **Table 1**, GLYCAM-Web hosts a JSON-API that supports interoperability with any other resource that can generate the GLYCAM condensed nomenclature from their representations, including GlyGen (glygen.org)^22^ and GlyConnect (glyconnect.expasy.org)^23^. Usage documentation for the JSON-API can be found on GitHub (github.com/GLYCAM-Web/website). The API facilitates the batch generation of structures, and the use of GLYCAM-Web within other cyber-infrastructure resources.

### Generating 3D structures from a carbohydrate sequence

The ability to generate reasonable 3D structures of carbohydrates (glycans, oligo- and polysaccharides) directly from their primary sequence results from their unique structural properties. At the monosaccharide level, pyranose rings generally strongly favor a single chair conformation due to 1,3 di-axial interactions^24^, simplifying their modeling. For furanoses, multiple, interconverting rings shapes complicate modeling, requiring MD simulations, but the impact of ring shape on the orientation of substituents is not profound^25^, meaning any single representative monosaccharide shape may be a reasonable initial structure for a simulation. For the great majority of monosaccharides available on GLYCAM-Web, only one ring shape is required for a representative 3D structure. At the oligosaccharide level the overall conformation of the polymer chain results from the values of the glycosidic dihedral angles, named φ, ψ, and ω, between the two monosaccharide residues (**Figure 3a**). As shown by experimentally-determined 3D structures, NMR observables, and quantum calculations, the φ angle of pyranose rings generally adopts only one low energy conformation dictated by the *exo*-anomeric effect (**Figure 3b**)^26^. Thus, a single, dominant φ conformation is, for the most part, adopted regardless of monosaccharide type or anomeric configuration. In fact, if the C2-C1-OX-CX definition is used, a φ value of approximately 180° is common for either anomeric configuration (α or β) (**Figure 3c**). In most two-bond linkages, the ψ angle also displays only one low energy conformation, attributed to steric effects^27^. In the case of three-bond linkages, such as at the O-6 position, the O6-C6-C5-O5 dihedral angle (ω) introduces additional flexibility, resulting in a preference for either two or three staggered rotamers (**Figure 3d**). Interestingly, the rotamer preference of the C6-C5 bond depends on the configuration of the C4 hydroxyl group. An equatorial OH group at C4, as in mannose and glucose, leads to 1-6 linkages that populate exclusively two of the three possible staggered rotamers (referred to as the *gg* and *gt* rotamers), whereas when the C4 hydroxyl group is axial, as in galactose, the ω-angle populates all three rotamers (*gg, gt, tg*). These preferences can be attributed to electrostatic repulsions between the O6 and O4 atoms (**Figure 3d**)^4,28^. For these reasons, and in contrast to oligopeptides, oligosaccharides adopt a relatively small set of well-defined conformations, enabling their 3D shapes to be readily predicted^29,30^.

**Figure 3.**
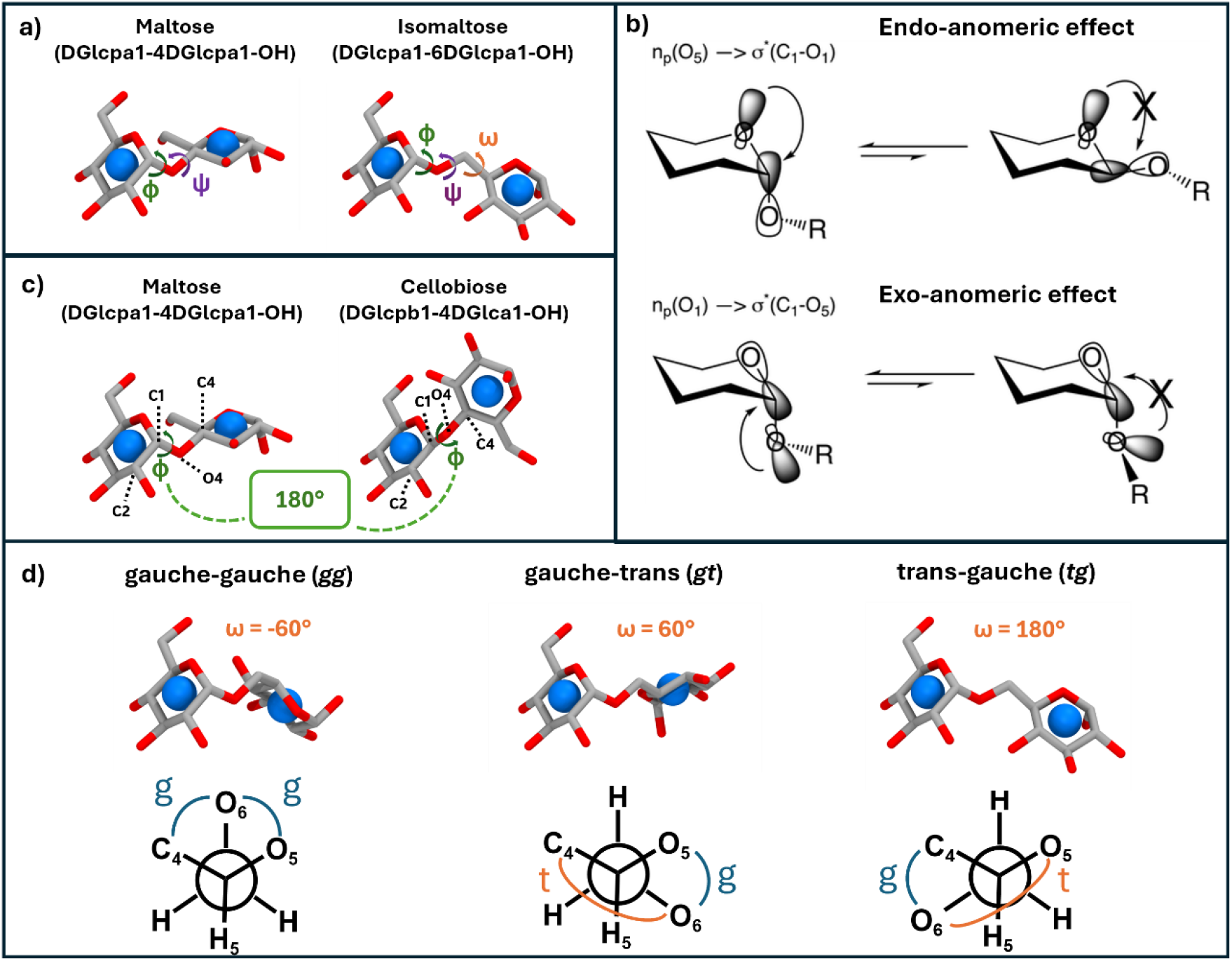
Glycosidic linkages adopt a small set of well-defined conformations. a) Carbohydrates are commonly linked together via two- or three-bond linkages, where the conformation can be defined by φ, ψ, and ω dihedral angles. b) The preferred orientation of the φ angle is dictated by the *exo*-anomeric effect; reprinted with permission^29^. c) Defining the φ dihedral angle using the C2-C1-Ox-Cx atoms results in an *exo*-anomeric value of approximately 180° for either α or β linkages. d) In contrast to the φ and ψ angles, the ω angle can exhibit up to three minima depending on the monosaccharide on the reducing end of the linkage. Isomaltose (DGlcpa1-6DGlcpa1-OH) is used here as an example. Carbohydrates shown in licorice format with 3D-SNFG symbols^31^.

### The carbohydrate sequence nomenclature used on GLYCAM-Web

Numerous nomenclatures for carbohydrates have been proposed that range widely in complexity and generalizability, depending on the initial motivation for their creation. The GLYCAM condensed nomenclature (similar to the subsequent IUPAC condensed nomenclature^32^) was adopted in order to facilitate the description and modeling of an almost unlimited number of complex carbohydrates. Over time, nomenclatures have not converged, but rather, software has been produced that enables their interconversion^33^. Regardless of the nomenclature, to generate a 3D structure, all sequences must be fully defined without ambiguity. The anomericity (α/β, simplified to a/b), ring form (furanose, f, or pyranose, p) and isomer (D/L) must be specified. The nature and configuration of the reducing terminus must also be specified, for example as α-OH, β-OMe, etc (See examples in **Table 2**).

**Table 2:**
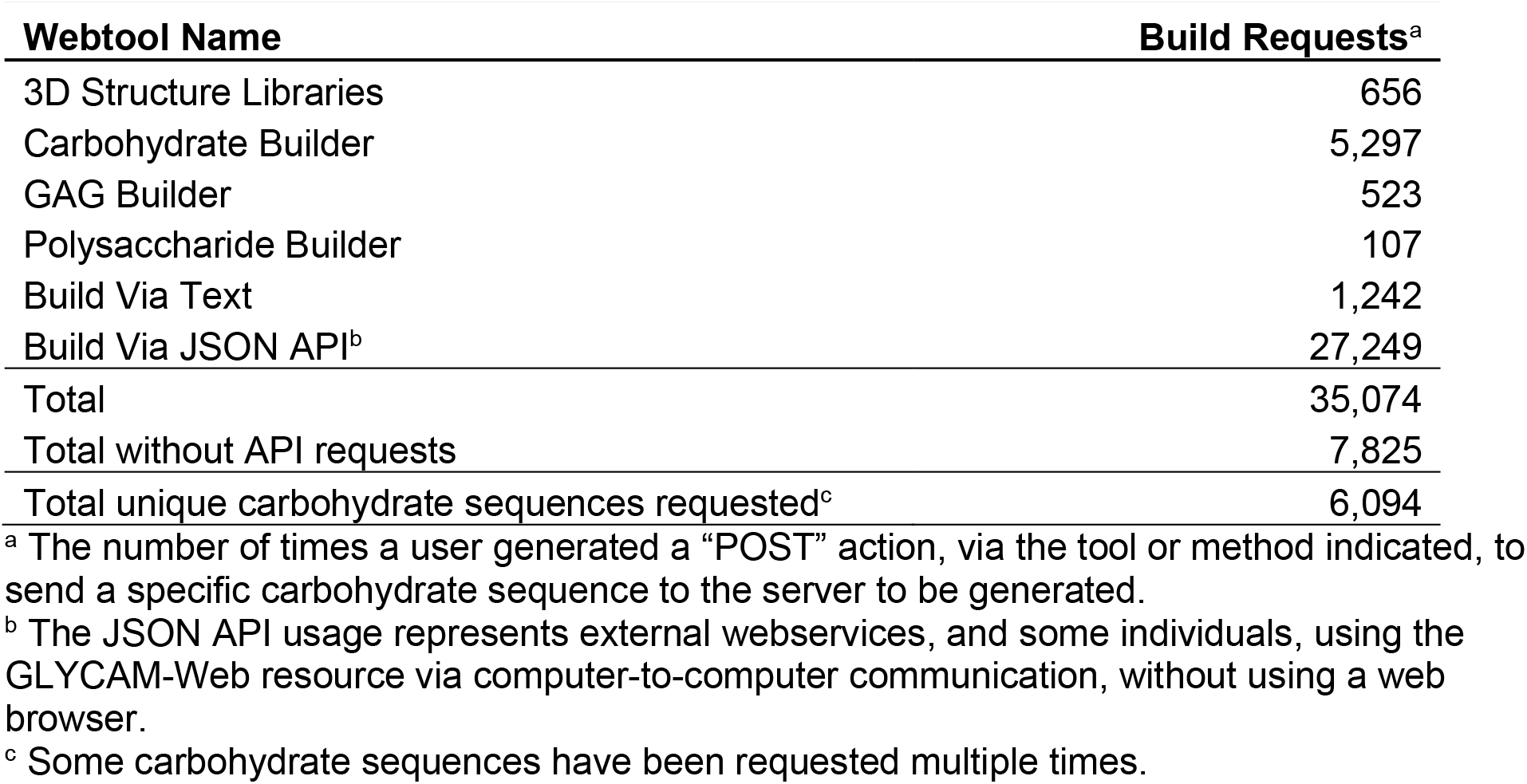
One-year usage data for GLYCAM-Web carbohydrate builders (2023 - 2024).

The GLYCAM condensed nomenclature supports branched oligosaccharides, in which the branches are contained within square brackets, for example the 1-6 branch in the trimannoside core of an *N*-glycan: DManpa1-3[DManpa1-6]DManpb1-4DGlcpNAcb1-4DGlcpNAcb1-OH. In this sequence, the DManpb residue is substituted at both the 3 and 6 positions, with the 1-6 linkage arbitrarily chosen as the branch point, rather than the 1-3 linkage, and included within square brackets. The alternate definition (DManpa1-6[DManpa1-3]DManpb1-4DGlcpNAcb1-4DGlcpNAcb1-OH), with the 1-3 linkage in square brackets, would also result in the same 3D structure. The degree of branching is unlimited within the nomenclature. Derivatives such as sulfate (S), methyl (Me), acetyl (Ac), are also specified within square brackets, placed between the ring form and anomer. For example, a galactopyranose substituted at the 2, 3, and 4 positions with Ac, S, and Me, respectively, would be represented as: DGalp[4Ac,2S,3Me]b1-OH. Deoxy positions are indicated in the similarly. For example, 6-deoxy-L-galactopyranose would be written as: LGalp[6H]b1-OH. However, because 6-deoxy-L-galactopyranose is a common monosaccharide (L-fucose), it has its own name: LFucpb1-OH. Large polymers may also be specified using square brackets to enclose the polymeric repeating unit, with the number of repeats subsequently specified within angled brackets (**Table 2**). As noted above, carbohydrate polymers need not be linear, as illustrated with the following 25-mer sequence for the capsular polysaccharide from group B *streptococcus* (serotype III)^34^: [4DGlcpb1-6[DNeup5Aca2-3DGalpb1-4]DGlcpNAcb1-3DGalpb1-]<25>OH, which highlights the convenience of the GLYCAM condensed nomenclature for representing highly complex carbohydrate structures. Non-reducing linkages between monosaccharides, such as in sucrose, are supported as follows: DFrufb2-1DGlcpa. In the special case of iduronate residues, which are known to populate multiple ring forms^35–38^, the ring pucker (^1^C_4_ (1C4), ^2^S_O_ (2SO), ^4^C_1_ (4C1)) can also be specified inside parentheses preceding any derivatives.

As polymeric sequences may also be heterogeneous, head and tail sequences may be defined around a repeating unit, for example the sequence for a 10-mer polymer of a heparan sulfate fragment that includes the biologically-important tail sequence would be defined as: [4DGlcpAb1-4DGlcpNAca1-]<10>4DGlcpAb1-3DGalpb1-3DGalpb1-4DXylpb1-OH (see **Table 2**).

### Accuracy and experimental validation

The 3D structure and conformational preferences of the monosaccharides available on GLYCAM-Web have been established (see Methods). At the oligosaccharide level, both NMR data and repositories of experimentally determined 3D structures can be used to derive appropriate dihedral angles for glycosidic linkages^29,39–46^. It is important to note that determining a single 3D structure for a flexible molecule, such as a carbohydrate, from NMR data can lead to the generation of a “virtual conformation”^47^. However, MD simulations of carbohydrates in water may be employed to identify dominant conformational states, which can be employed to deconvolute NMR observables, such as scalar three-bond couplings (^3^*J*), into experimentally-consistent populations for each of the solution conformations^48^.

As glycan-binding proteins most often recognize the predominant solution conformation, the data from 3D structure databases can also be integrated to determine what states are possible and how frequently they are populated^29,30,39,49^. The carbohydrate-containing structures (>14,000^50^ instances) in the Protein Data Bank^51^ comprise an obvious validation dataset, but caution must be employed as a lack of standards and carbohydrate-specific biocuration tools during deposition has allowed for numerous forms of errors in the experimental data^52^. To generate a high-quality dataset from the PDB, only carbohydrates with average b-factors of less than 30 Å^2^ were selected (using the GlyFinder tool, www.glycam.org/gf) for comparison to the predicted shapes. Further, the PDB data were filtered to remove any highly distorted ring shapes, and biologically inconsistent inter-residue linkages. For example, an *N*-linked glycan containing a reported Man-α1-4-GlcNAc linkage would be excluded, as the linkage must be β1-4. A comparison of the energy minimized structures from the carbohydrate builder to this high-quality data to is shown in Figure 4. The vast majority of β-linkages in the PDB were β1-4 linkages (n=3229), and the average shape, as measured by the φ and ψ dihedral angles of the linkages, was well reproduced by the carbohydrate builder structure (φ Z-score = 0.27, ψ Z-score = 0.49). While fewer examples of β1-2 linkages are present in the PDB, the carbohydrate builder structures are in agreement with the experimental data. In the case of β1-3 linkages, there were too few examples (n=3) to draw statistically significant conclusions. The carbohydrate builder also performed well for α-linkages, reproducing the two distinct shapes adopted by the ψ angle of various α1-2 linkages, as well as the single shapes adopted by the other linkages.

**Figure 4:**
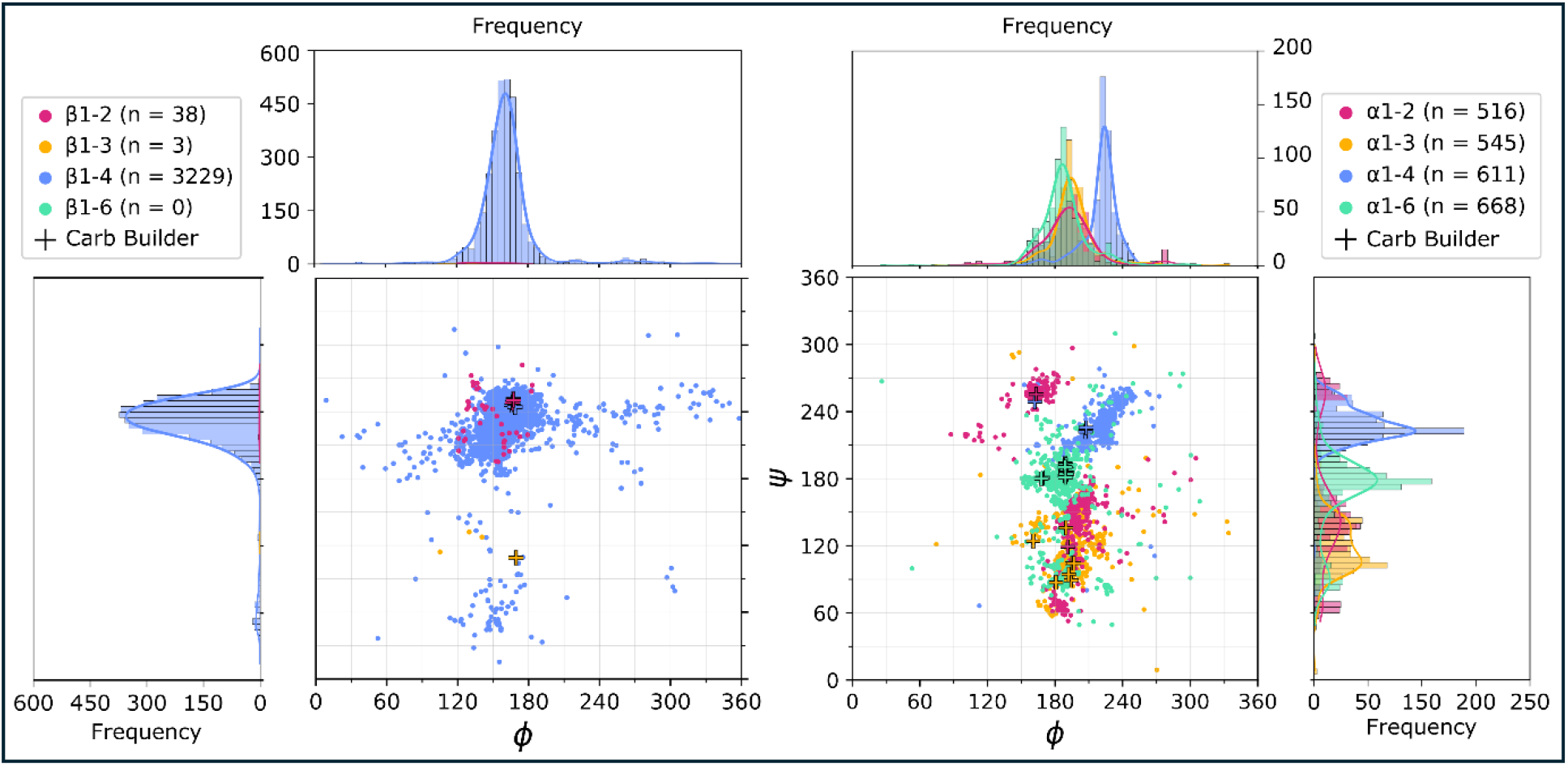
A comparison of the dihedral angle distributions for the glycosidic linkages generated by the GLYCAM-Web carbohydrate builder and high-quality experimentally determined structures in the PDB. PDB data were retrieved using the GlyFinder webtool (glycam.org/gf) and filtered as described. Frequency histograms of the φ and ψ dihedral angles showed distinct preferences between linkage types. These preferences were reproduced by the energy minimized structures generated by the carbohydrate builder. Dihedral angle definitions: φ: C_2_-C_1_-O_X_-C_X_, ψ: C_1_-O_X_-C_X_-C_X+1_ for 1-2, 1-3 and 1-4 linkages; φ: C_2_-C_1_-O_6_-C_6_, ψ: C_1_-O_6_-C_6_-C_5_ for 1-6 linkages.

## Methods

### Sequence parsing and initial structure generation

The various carbohydrate builders presented here ultimately send a text-format sequence (e.g. DManpa1-2DGlcpb1-OH) to underlying scientific software via a JSON API. The open-source software currently employed by GLYCAM-Web, called the GLYCAM Molecular Modeling Library (GMML), was developed in the Woods group, is written in C++ and contains high-level features for the generation and manipulation of carbohydrate 3D structures. It is publicly available via GitHub (https://github.com/GLYCAM-Web/gmml2) and when installed can be used to compile the carbohydrate builder program as a stand-alone executable. The carbohydrate builder requires a carbohydrate sequence in GLYCAM condensed nomenclature, which is converted into a graph structure of connected monosaccharides, and a lookup table is used to find the appropriate 3D monosaccharide template files taken from the GLYCAM06 force field^53^, which contain the atom names, Cartesian coordinates, atom types, partial atomic charges and atomic connectivities. The 3D structures and default ring forms for each monosaccharide have been taken from the available literature (**Table 3**). The oligosaccharide is then assembled sequentially from the constituent monosaccharides according to the known preferences of each glycosidic linkage.

**Table 3:**
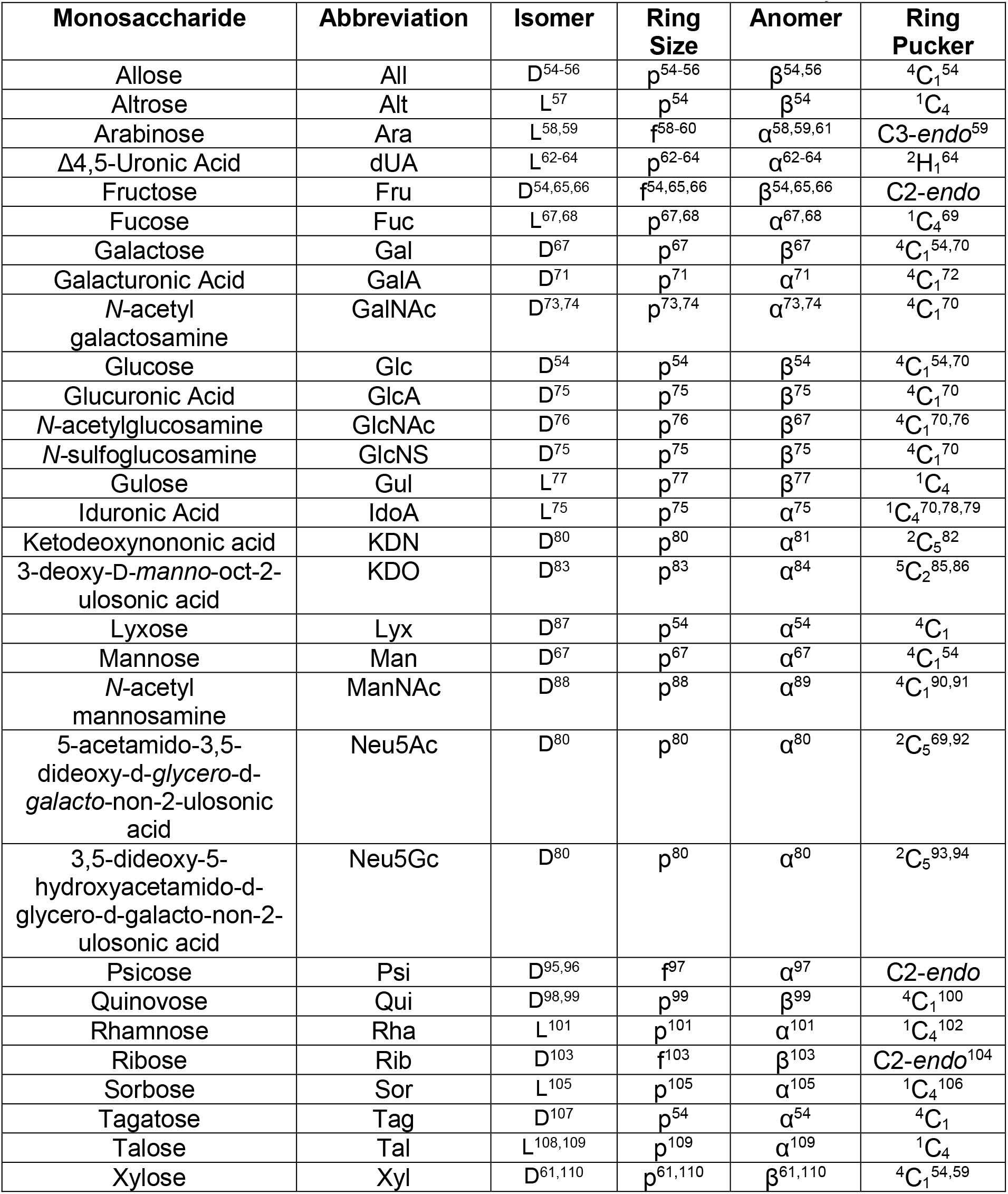
Default features for monosaccharides in the GLYCAM-Web carbohydrate builders.

### Overlap resolution

The initial 3D structure is generated using experimentally consistent default values^41–45,53^ for the glycosidic bond distances and angles and dihedral angles (see **Table 4**). While this usually generates a viable structure, a unique shape or branching pattern of an oligosaccharide may lead to potential steric overlaps between residues, particularly in the case of large or non-natural oligosaccharides. Severe overlaps, although uncommon, can result in the energy minimization algorithm failing to find a reasonable 3D structure. To avoid this issue, the initial structures are assessed for the presence of atomic overlaps, which are resolved via a two-step process. Firstly, as each residue is added to the carbohydrate chain a “greedy”^111^ algorithm resolves any local atomic overlaps between the new residue and the existing residues by adjusting the dihedral angle value of the new glycosidic linkage within experimentally-defined bounds^49^. In the second stage, once the full structure is generated, any remaining overlaps are resolved by sequentially adjusting the dihedral angles of each glycosidic linkage. The default values for dihedral angles are taken from the literature where available, and more general rules are used when default values are not available (see **Table 4**). When adjusting for overlaps, the dihedral angles are currently explored in 1° increments to a maximum deviation of ± 20°, which was chosen as a conservative estimate of their range of motion^49^. Where multiple angles produce the same degree of overlap, the angle that is closest to the default value is used. Once the lowest possible overlap is found, the next dihedral angle is adjusted in the same manner. The resulting structure is termed the “default structure” and is returned to the user for visualization on the website while the structure is being simultaneously energy-minimized for visualization and download. In GMML, a non-bonded overlap is considered to exist if any interatomic distance is less than the sum of their vdW radii minus 0.6 Å.

**Table 4:**
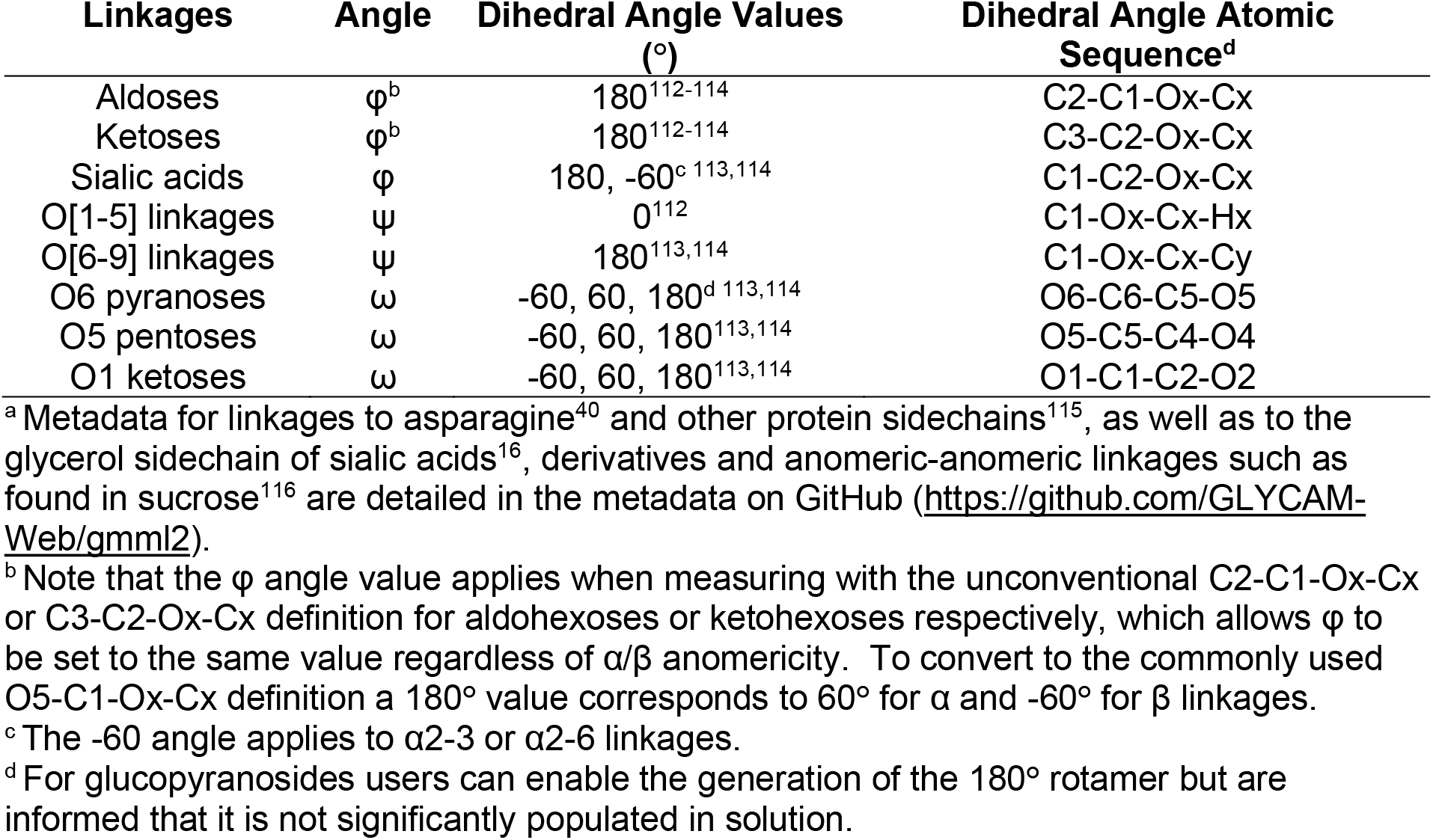
Default dihedral angle values for glycosidic linkages^a^.

| Linkages | Angle | Dihedral Angle Values (°) | Dihedral Angle Atomic Sequence<sup>d</sup> |
| --- | --- | --- | --- |
| Aldoses | $\phi^b$ | 180<sup>112-114</sup> | C2-C1-Ox-Cx |
| Ketoses | $\phi^b$ | 180<sup>112-114</sup> | C3-C2-Ox-Cx |
| Sialic acids | $\phi$ | 180, -60<sup>c 113,114</sup> | C1-C2-Ox-Cx |
| O[1-5] linkages | $\psi$ | 0<sup>112</sup> | C1-Ox-Cx-Hx |
| O[6-9] linkages | $\psi$ | 180<sup>113,114</sup> | C1-Ox-Cx-Cy |
| O6 pyranoses | $\omega$ | -60, 60, 180<sup>d 113,114</sup> | O6-C6-C5-O5 |
| O5 pentoses | $\omega$ | -60, 60, 180<sup>113,114</sup> | O5-C5-C4-O4 |
| O1 ketoses | $\omega$ | -60, 60, 180<sup>113,114</sup> | O1-C1-C2-O2 |

<sup>a</sup> Metadata for linkages to asparagine<sup>40</sup> and other protein sidechains<sup>115</sup>, as well as to the glycerol sidechain of sialic acids<sup>16</sup>, derivatives and anomeric-anomeric linkages such as found in sucrose<sup>116</sup> are detailed in the metadata on GitHub (https://github.com/GLYCAM-Web/gmml2).
<sup>b</sup> Note that the $\phi$ angle value applies when measuring with the unconventional C2-C1-Ox-Cx or C3-C2-Ox-Cx definition for aldohexoses or ketohexoses respectively, which allows $\phi$ to be set to the same value regardless of $\alpha/\beta$ anomericity. To convert to the commonly used O5-C1-Ox-Cx definition a 180° value corresponds to 60° for $\alpha$ and -60° for $\beta$ linkages.
<sup>c</sup> The -60 angle applies to $\alpha$ 2-3 or $\alpha$ 2-6 linkages.
<sup>d</sup> For glucopyranosides users can enable the generation of the 180° rotamer but are informed that it is not significantly populated in solution.

### Energy minimization protocol and output

The simulation protocols in use by GLYCAM-Web are made available on GitHub (https://github.com/GLYCAM-Web/MD_Utils.git). GMML outputs an overlap resolved “default structure” as an AMBER format OFF file. The tleap module of AmberTools^117^ is subsequently used to generate AMBER format PRMTOP and INPCRD files, which are then used to perform an energy minimization with a dielectric constant appropriate to water. The resulting structure is processed by cpptraj^118^ and then tleap^117^ to create explicitly solvated structures with either the TIP3P^119^ or TIP5P^120^ water models. These files are then made available for download.

### Handling derivatives including deoxy positions

GLYCAM residues were designed to be modular and transferable while retaining reasonable electrostatic properties. To this end, charges at linking atoms are adjusted so as to maintain overall integral net molecular charge^53^. The partial charge adjustments performed for each of the supported derivatives are detailed in **Table 5**. To create a deoxy position in a monosaccharide, the O and H atoms are deleted and replaced with an H atom. The charges from the deleted atoms are added to the atomic charge of the deoxy C atom. This protocol is not exactly equivalent to that which would be used to compute the partial atomic charges denovo for such a deoxy monosaccharide^8^, although the resulting charges are similar. Thus, the atomic charges for the common deoxy monosaccharide LFucp will not be identical to those for the equivalent monosaccharide derived from 6-deoxy L-Gal (LGal[6H]p). For this reason, users are recommended to select common deoxy monosaccharides if they are present in the carbohydrate builder.

**Table 5:** Charge adjustment performed by GLYCAM-Web after derivatization of glycan <u>hydroxyl groups.</u>.

| Derivatization Type | GLYCAM Residue Code | Charge Adjustment Atom | Charge adjustment |
| --- | --- | --- | --- |
| Acetylation | ACX | C-O-Derivative<sup>a</sup> | +0.008 |
| Methylation | MEX | C-O-Derivative<sup>a</sup> | -0.039 |
| Sulfation | SO3 | C-O-Derivative<sup>b</sup> | +0.031 |

<sup>a</sup> The charge adjustment is applied to the carbon atom bonded to the linking atom. For example, if O3 is acetylated or methylated the charge for the C3 atoms is adjusted.
<sup>b</sup> The charge adjustment is applied to the derivatized oxygen.

In some cases, monosaccharides exhibit different features depending on their presence within an oligosaccharide or as a monosaccharide in solution. For example, in the monosaccharide form, fructose is most stable as β-D-fructopyranose^121^, however the furanose form is dominant in plant oligosaccharides like raffinose^122^ and sucrose^123^. Thus, when selecting default monosaccharide features, priority was given to the dominant form as it occurs in oligosaccharides. If sufficient literature was not available to indicate the dominant form, then the features for the single monosaccharide in solution were selected. In the absence of reported data, the default features were chosen based on related monosaccharides. A single ring pucker was selected for pyranoses, either ^4^C_1_ or ^1^C_4_, depending on if the pyranose was the D or L isomer, respectfully. Similarly, for furanoses, a single pucker of C2-*endo* and C3-*endo*, respectively. Note that for furanoses, the energy barrier is very small between the C2-*endo* and C3-*endo* puckers, so the minimized pucker may differ from the initial pucker that is assigned^25^. Sialic acid puckers are equivalent to those of the other pyranoses, but the ring atom numbering is different. In sialic acids, C2 is the anomeric carbon, so ^2^C_5_ and ^5^C_2_ are equivalent to ^1^C_4_ and ^4^C_1_, respectively. However, in general, the default puckers are not arbitrary, and references are provided where possible. L-IdoA is mainly found in Heparin/Heparan Sulfate glycosaminoglycans and exhibits a ring puckering equilibrium between ^1^C_4_ and ^2^S_O_. This equilibrium is dependent on many factors, including neighboring sulfation and protein binding. ^1^C_4_ was selected as the default.

## Discussion

### Community Adoption

A webtool has several innate advantages over standalone software, including that no software needs to be installed or updated by the user, reducing barriers to adoption, especially by non-experts. Further, the automation required for a web-based implementation necessitates the adoption of standardized protocols. This reproducibility combined with robust operation and ease of use have contributed to the widespread adoption of the tools at GLYCAM-Web (**Figure 5, Table 6**).

**Figure 5.**
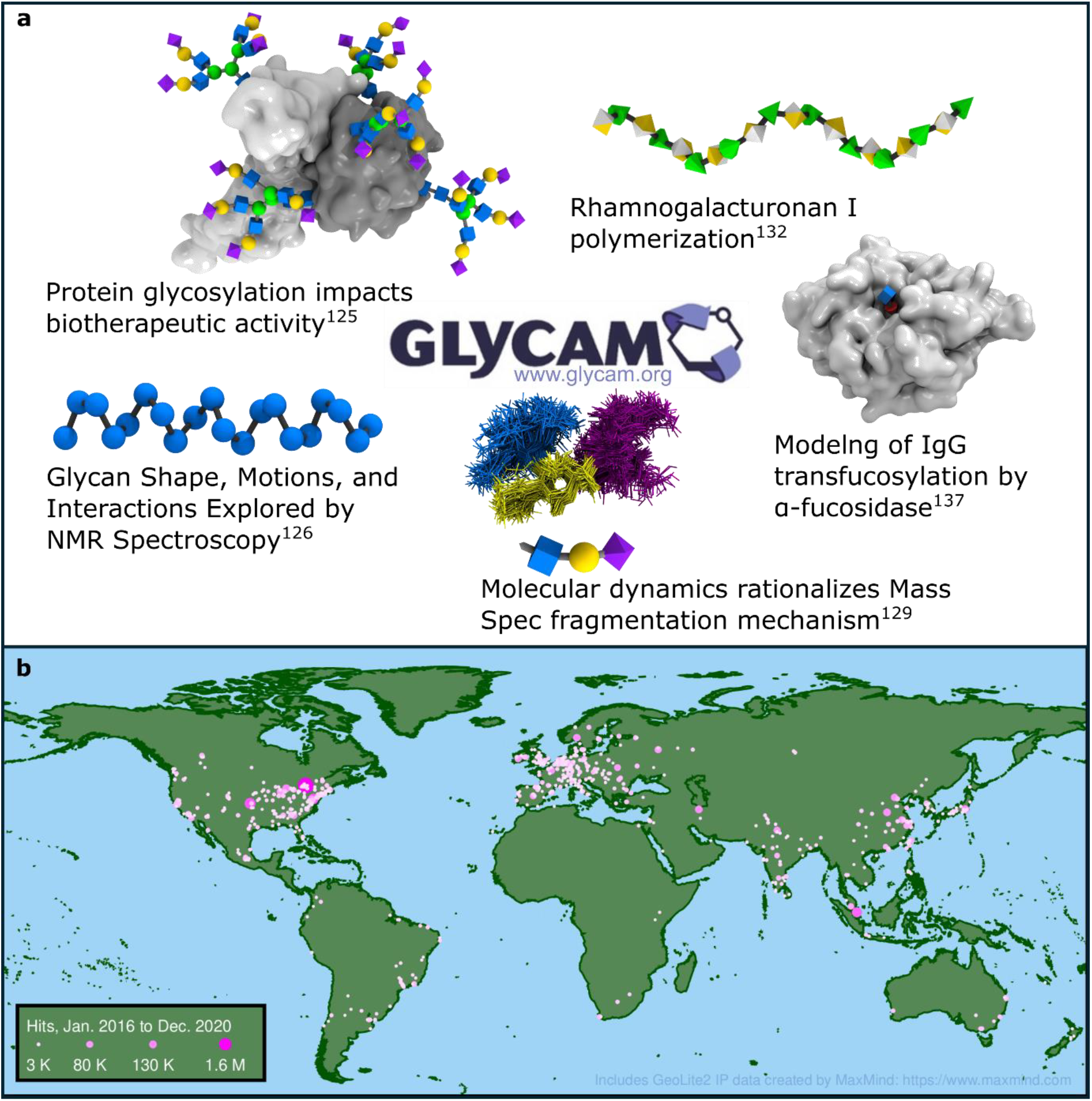
a) A representation of the broad range of research enabled by the GLYCAM-Web carbohydrate builders (the examples represent research cited in **Table 7**). b) A plot of global website hits spanning Jan 2016 – Dec 2020. Access from web-crawlers and our developers has been omitted as much as possible. A hit is defined as any single request made to the webserver by a client. For clarity any location with less than 1000 hits is excluded.

The data in **Figure 5** corresponds to approximately 372,698 logged visits over a 5-year period, growing to 80,000 visits per year in 2020 (due to system changes visitor statistics were not recorded from 2021-2024). Focusing on recent usage of the carbohydrate builders (**Table 2**) it indicates that approximately 21 build requests were processed per day. This figure ignores the large number of website-to-website external builds employing the JSON API.

A google scholar search for publications referencing the terms “GLYCAM-Web” or the GLYCAM-Web URL (www.glycam.org) identified approximately 350 publications from 2020 through 2024 (see Supplementary Data). An indication of the breadth and depth of the impact of GLYCAM-Web, driven in large part by use of the carbohydrate builders presented herein, is provided by the selection of articles listed in **Table 7**.

**Table 7:** Fifteen representative peer-reviewed articles from 2020-2024 citing the use of GLYCAM-Web.

| Article Title | Journal | Ref. |
| --- | --- | --- |
| A consensus structural motif for the capsular polysaccharide of *Cryptococcus Neoformans* by NMR/MD | Proc. Natl. Acad. Sci., USA | 124 |
| Exposing the molecular heterogeneity of glycosylated biotherapeutics | Nat. Commun. | 125 |
| The essential malaria protein PfCyRPA targets glycans to invade erythrocytes | Cell Rep. | 126 |
| Restoring protein glycosylation with GlycoShape | Nat. Methods | 127 |
| Small lectin ligands as a basis for applications in glycoscience and glycomedicine | Chem. Soc. Rev. | 128 |
| Predicting glycan structure from tandem mass spectrometry via deep learning | Nat. Methods | 129 |
| Glycan Shape, Motions, and Interactions Explored by NMR Spectroscopy | J. Am. Chem. Soc., Au | 130 |
| SARS-CoV-2 uses CD4 to infect T helper lymphocytes | Elife | 131 |
| Polymerization of the backbone of the pectic polysaccharide rhamnogalacturonan I | Nat. Plants | 132 |
| Mechanism of mixed-linkage glucan biosynthesis by barley cellulose synthase-like CslF6 (1, 3; 1, 4)-β-glucan synthase | Sci. Adv. | 133 |
| SARS-CoV-2 spike opening dynamics and energetics reveal the individual roles of glycans and their collective impact | Commun. Biol. | 134 |
| Revealing the evolution towards complex *N*-glycan specificities of human H1 influenza A viruses | J. Am. Chem. Soc., Au | 135 |
| NMR of glycoproteins: profiling, structure, conformation and interactions | Curr. Opin. Struct. Biol. | 136 |
| Structure and dynamics of an α-fucosidase reveal a mechanism for highly efficient IgG transfucosylation | Nat. Commun | 137 |
| Insight into the catalytic mechanism of GH11 xylanase: computational analysis of substrate distortion based on a neutron structure | J. Am. Chem. Soc. | 138 |

### Future directions

After 20 years of operation, the GLYCAM-Web suite webtools have evolved as a result of widespread community adoption and funding mechanisms that support cyberinfrastructure, and a dedicated team of professionals that handle cybersecurity, web development, system and network administration, development operations, as well as software development and maintenance.

As noted in the introduction, numerous glycan nomenclatures exist, as do tools for their interconversion^33^. To further advance the interoperability of the GLYCAM-Web tools, it would be helpful to export the GLYCAM Condensed nomenclature in other formats, such as WURCS^139^ or the similar IUPAC Condensed^32,140^. Support for additional file formats, such as the crystallization Information File (CIF)^141^ for 3D structures would help to facilitate the analysis of cryo-EM or X-Ray diffraction data.

MD simulations are highly effective in modeling the flexibility of carbohydrates, yet their execution requires expertise in biomolecular simulations as well as carbohydrate chemistry, which introduces a significant bottleneck to their widespread application. To surmount this difficulty, an online MD service is in beta-testing at GLYCAM-Web. This service enables registered users to submit 3D structures created with the carbohydrate builders at GLYCAM-Web for online MD simulation. This capability provides the research community with a set of expert-curated protocols and standardized, reproducible MD simulations of carbohydrates in solution. In future, this service could be enhanced by providing automated analyses of the resulting simulation data, for example by automating the calculation of NMR properties, structural features, etc., for direct comparison with experimental data. Increasing the number of chemical or biological derivatives and expanding the repertoire of monosaccharides available to include, for example, bacterial monosaccharides would further broaden the application space of the carbohydrate builders.

## Code availability

The GEMS and GMML2 software repositories are open-source (GNU Lesser General Public License v3.0) and publicly accessible on GitHub (https://github.com/GLYCAM-Web). For security, the website code and deployment pipeline have not been made public, however the architecture currently in place has been designed to be transferable to perpetuate its use.

## Acknowledgments

GLYCAM-Web was directly supported by NIH R24GM136984 (Transitioning Glycam-Web to a self-sustaining carbohydrate modeling service), U01CA207824 (Tools to enable non-specialists) and R01GM100058 (Continued Development and Maintenance of GLYCAM-Web). This work is currently supported by NIAID R01AI155975 and GlycoMIP, a National Science Foundation Materials Innovation Platform funded through Cooperative Agreement DMR-1933525. David Sehnal acknowledges funding from the Grant Agency of Czech Republic JuniorStar project (22-30571M). Xiaocong Wang is supported by Hubei Provincial Natural Science Foundation of China, No. 2025AFB695.

