## Supplemental Information for "Generating 3D Models of Carbohydrates with GLYCAM-Web"

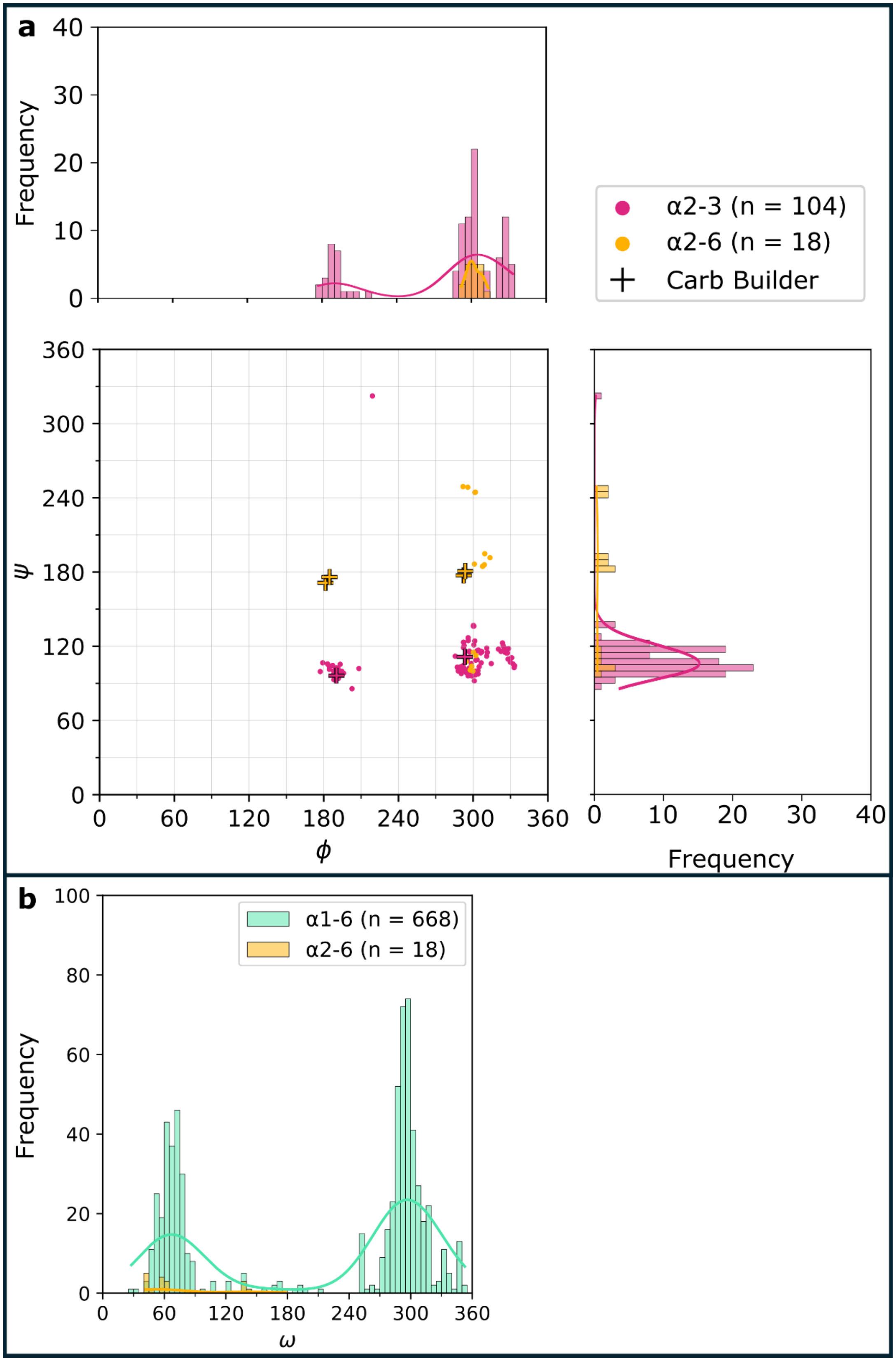
a
b
Frequency
α2-3 (n = 104)
α2-6 (n = 18)
Carb Builder
ψ
ϕ
Frequency
α1-6 (n = 668)
α2-6 (n = 18)
Frequency
ω

**Supplemental Figure S1**: A comparison of the dihedral angle distributions for the glycosidic linkages generated by the GLYCAM-Web carbohydrate builder and high-quality experimentally determined structures in the PDB. PDB data were retrieved using the GlyFinder webtool (glycam.org/gf) and filtered as described. a) A plot of φ versus ψ as well as frequency histograms for α2-3 and α2-6 linkages. b) A frequency histogram for the ω angle of α1-6 and α2-6 linkages. The values for carbohydrate builder structures for the ω angle are 60 (gt), 180 (tg) and 300 (gg). Dihedral definitions φ: $C_1$-$C_2$-$O_3$-$C_3$, ψ: $C_2$-$O_3$-$C_3$-$C_4$ for 2-3 linkage, φ: $C_1$-$C_2$-$O_6$-$C_6$, ψ: $C_2$-$O_6$-$C_6$-$C_5$ for 2-6 linkage, ω: $O_6$-$C_6$-$C_5$-$O_5$.

**Supplementation references**: manuscripts published in 2020[1-69], 2021[70-141], 2022[142-221], 2023[222-279] and 2024[280-349] that acknowledge or reference GLYCAM-Web (www.glycam.org) not authored by Dr. Robert J. Woods or members of his group.


1 Roe, D. R. & Brooks, B. R. A protocol for preparing explicitly solvated systems for stable molecular dynamics simulations. *J Chem Phys* **153**, 054123, https://doi.org/10.1063/5.0013849 (2020).

2 Bhattarai, S., Pippel, J., Scaletti, E., Idris, R., Freundlieb, M., Rolshoven, G., Renn, C., Lee, S. Y., Abdelrahman, A., Zimmermann, H., El-Tayeb, A., Muller, C. E. & Strater, N. 2-Substituted alpha,beta-Methylene-ADP Derivatives: Potent Competitive Ecto-5'-nucleotidase (CD73) Inhibitors with Variable Binding Modes. *J Med Chem* **63**, 2941-2957, https://doi.org/10.1021/acs.jmedchem.9b01611 (2020).

3 Tian, C., Kasavajhala, K., Belfon, K. A. A., Raguette, L., Huang, H., Migues, A. N., Bickel, J., Wang, Y., Pincay, J., Wu, Q. & Simmerling, C. ff19SB: Amino-Acid-Specific Protein Backbone Parameters Trained against Quantum Mechanics Energy Surfaces in Solution. *J Chem Theory Comput* **16**, 528-552, https://doi.org/10.1021/acs.jctc.9b00591 (2020).

4 Abrahams, J. L., Taherzadeh, G., Jarvas, G., Guttman, A., Zhou, Y. & Campbell, M. P. Recent advances in glycoinformatic platforms for glycomics and glycoproteomics. *Current opinion in structural biology* **62**, 56-69 (2020).

5 Chen, N., Gao, H.-X., He, Q., Yu, Z.-L. & Zeng, W.-C. Interaction and action mechanism of starch with different phenolic compounds. *International Journal of Food Sciences and Nutrition* **71**, 726-737 (2020).

6 Copoiu, L. & Malhotra, S. The current structural glycome landscape and emerging technologies. *Current Opinion in Structural Biology* **62**, 132-139 (2020).

7 Roy, R., Poddar, S. & Kar, P. Conformational preferences of triantennary and tetraantennary hybrid N-glycans in solution: Insights from 20 μs long atomistic molecular dynamic simulations. (2020).

8 Fogarty, C. A., Harbison, A. M., Dugdale, A. R. & Fadda, E. How and why plants and human N-glycans are different: Insight from molecular dynamics into the “glycoblocks” architecture of complex carbohydrates. *Beilstein journal of organic chemistry* **16**, 2046-2056 (2020).

9 Roy, R., Ghosh, B. & Kar, P. Investigating conformational dynamics of Lewis Y oligosaccharides and elucidating blood group dependency of cholera using molecular dynamics. *ACS omega* **5**, 3932-3942 (2020).

10 Zhang, W., Meredith, R. J., Oliver, A. G., Carmichael, I. & Serianni, A. S. Glycosidic linkage, N-acetyl side-chain, and other structural properties of methyl 2-acetamido-2-deoxy-β-d-glucopyranosyl-(1→ 4)-β-d-mannopyranoside monohydrate and related compounds. *Acta Cryst.* **76**, 287-297 (2020).

11 Lukose, B. & Rani, P. G82S RAGE polymorphism influences amyloid-RAGE interactions relevant in Alzheimer’s disease pathology. *PLoS One* **15**, e0225487 (2020).

12 Chen, N., Chen, L., Gao, H. X. & Zeng, W. C. Mechanism of bridging and interfering effects of tea polyphenols on starch molecules. *Journal of Food Processing and Preservation* **44**, e14576 (2020).
13 Uslupehlivan, M. & Şener, E. Computational analysis of SARS-CoV-2 S1 protein O-glycosylation and phosphorylation modifications and identifying potential target positions against CD209L-mannose interaction to inhibit initial binding of the virus. *bioRxiv*, 2020.2003. 2025.007898 (2020).
14 Cavada, B. S., Osterne, V. J. S., Pinto-Junior, V. R., Souza, L. A. G., Lossio, C. F., Silva, M. T. L., Correia-Neto, C., Oliveira, M. V., Correia, J. L. A. & Neco, A. H. B. Molecular dynamics and binding energy analysis of Vatairea guianensis lectin: a new tool for cancer studies. *Journal of Molecular Modeling* **26**, 1-9 (2020).
15 de Oliveira Leite, G., Santos, S. A. A. R., Bezerra, F. M. D. H., e Silva, F. E. S., de Castro Ribeiro, A. D., Roma, R. R., Silva, R. R. S., Santos, M. H. C., Santos, A. L. E. & Teixeira, C. S. Is the orofacial antinociceptive effect of lectins intrinsically related to their specificity to monosaccharides? *International Journal of Biological Macromolecules* **161**, 1079-1085 (2020).
16 Lal, K., Bermeo, R. & Perez, S. Computational tools for drawing, building and displaying carbohydrates: a visual guide. *Beilstein Journal of Organic Chemistry* **16**, 2448-2468 (2020).
17 Re, S., Yamaguchi, Y. & Sugita, Y. Molecular dynamics simulation of glycans. *Trends in Glycoscience and Glycotechnology* **32**, E113-E118 (2020).
18 Scherbinina, S. I. & Toukach, P. V. Three-dimensional structures of carbohydrates and where to find them. *International journal of molecular sciences* **21**, 7702 (2020).
19 Barnett, C. B., Senapathi, T. & Naidoo, K. J. Comparative ligand structural analytics illustrated on variably glycosylated MUC1 antigen–antibody binding. *Beilstein Journal of Organic Chemistry* **16**, 2540-2550 (2020).
20 Kong, R., Liu, R. R., Xu, X. M., Zhang, D. W., Xu, X. S., Shi, H. & Chang, S. Template-based modeling and ab-initio docking using CoDock in CAPRI. *Proteins: Structure, Function, and Bioinformatics* **88**, 1100-1109 (2020).
21 Zhou, Y., Zhao, Y., Niu, B., Luo, Q., Zhang, Y., Quan, G., Pan, X. & Wu, C. Cyclodextrin-based metal-organic frameworks for pulmonary delivery of curcumin with improved solubility and fine aerodynamic performance. *Int J Pharm X* **588**, 119777 (2020).
22 Plamitzer, L. & Bouř, P. Pressure dependence of vibrational optical activity of model biomolecules. A computational study. *Chirality* **32**, 710-721 (2020).
23 Shivgan, A. T., Marzinek, J. K., Huber, R. G., Krah, A., Henchman, R. H., Matsudaira, P., Verma, C. S. & Bond, P. J. Extending the Martini coarse-grained force field to N-glycans. *Journal of Chemical Information and Modeling* **60**, 3864-3883 (2020).
24 Martínez, J. D., Infantino, A. S., Valverde, P., Diercks, T., Delgado, S., Reichardt, N.-C., Ardá, A., Cañada, F. J., Oscarson, S. & Jimenez-Barbero, J. The interaction of fluorinated glycomimetics with DC-SIGN: multiple binding modes disentangled by the combination of NMR methods and MD simulations. *Pharmaceuticals* **13**, 179 (2020).
25 Wu, K., Li, D., Xiu, P., Ji, B. & Diao, J. O-GlcNAcylation inhibits the oligomerization of alpha-synuclein by declining intermolecular hydrogen bonds through a steric effect. *Phys. Biol.* **18**, 016002 (2020).
26 Srivastava, A. D., Unione, L., Wolfert, M. A., Valverde, P., Ardá, A., Jiménez-Barbero, J. & Boons, G. J. Mono-and Di-Fucosylated Glycans of the Parasitic Worm S. Mansoni Are Recognized Differently by the Innate Immune Receptor DC-SIGN. *Chemistry–A European Journal* **26**, 15605-15612 (2020).
27 Soliman, C., Guy, A. J., Chua, J. X., Vankemmelbeke, M., McIntosh, R. S., Eastwood, S., Truong, V. K., Elbourne, A., Spendlove, I. & Durrant, L. G. Molecular and structural basis

for Lewis glycan recognition by a cancer-targeting antibody. *Biochemical Journal* **477**, 3219-3235 (2020).
28 Kanampalliwar, A. & Singh, D. V. Extracellular DNA builds and interacts with vibrio polysaccharide in the biofilm matrix formed by Vibrio cholerae. *Environmental Microbiology Reports* **12**, 594-606 (2020).
29 Marchiori, M. F., Bortot, L. O., Carvalho, I. & Campo, V. L. Synthesis of MUC1-derived glycopeptide bearing a novel triazole STn analog. *Carbohydrate Research* **498**, 108155 (2020).
30 Stratilová, B., Šesták, S., Mravec, J., Garajová, S., Pakanová, Z., Vadinová, K., Kučerová, D., Kozmon, S., Schwerdt, J. G. & Shirley, N. Another building block in the plant cell wall: Barley xyloglucan xyloglucosyl transferases link covalently xyloglucan and anionic oligosaccharides derived from pectin. *The Plant Journal* **104**, 752-767 (2020).
31 Manjon, E., Bras, N. F., Garcia-Estevez, I. & Escribano-Bailón, M. T. Cell wall mannoproteins from yeast affect salivary protein–flavanol interactions through different molecular mechanisms. *Journal of Agricultural and Food Chemistry* **68**, 13459-13468 (2020).
32 Klontz, E. H., Li, C., Kihn, K., Fields, J. K., Beckett, D., Snyder, G. A., Wintrode, P. L., Deredge, D., Wang, L.-X. & Sundberg, E. J. Structure and dynamics of an α-fucosidase reveal a mechanism for highly efficient IgG transfucosylation. *Nature Communications* **11**, 6204 (2020).
33 Su, L., Yao, K. & Wu, J. Improved activity of sulfolobus acidocaldarius maltooligosyltrehalose synthase through directed evolution. *Journal of agricultural and food chemistry* **68**, 4456-4463 (2020).
34 Thompson, A. J., Cao, L., Ma, Y., Wang, X., Diedrich, J. K., Kikuchi, C., Willis, S., Worth, C., McBride, R. & Yates, J. R. Human influenza virus hemagglutinins contain conserved oligomannose N-linked glycans allowing potent neutralization by lectins. *Cell Host & Microbe* **27**, 725-735. e725 (2020).
35 Hembach, L., Bonin, M., Gorzelanny, C. & Moerschbacher, B. M. Unique subsite specificity and potential natural function of a chitosan deacetylase from the human pathogen Cryptococcus neoformans. *Proceedings of the National Academy of Sciences* **117**, 3551-3559 (2020).
36 Patro, L. P. P., Sudhakar, K. U. & Rathinavelan, T. K-PAM: a unified platform to distinguish Klebsiella species K-and O-antigen types, model antigen structures and identify hypervirulent strains. *Scientific Reports* **10**, 16732 (2020).
37 Jayaprakash, N. G., Singh, A., Vivek, R., Yadav, S., Pathak, S., Trivedi, J., Jayaraman, N., Nandi, D., Mitra, D. & Surolia, A. The barley lectin, horcolin, binds high-mannose glycans in a multivalent fashion, enabling high-affinity, specific inhibition of cellular HIV infection. *Journal of Biological Chemistry* **295**, 12111-12129 (2020).
38 Mishra, N., Sharma, S., Dobhal, A., Kumar, S., Chawla, H., Singh, R., Das, B. K., Kabra, S. K., Lodha, R. & Luthra, K. A rare mutation in an infant-derived HIV-1 envelope glycoprotein alters interprotomer stability and susceptibility to broadly neutralizing antibodies targeting the trimer apex. *Journal of Virology* **94**, 10.1128/jvi. 00814-00820 (2020).
39 Dussouy, C., Téletchéa, S., Lambert, A., Charlier, C., Botez, I., Ceuninck, F. d. r. D. & Grandjean, C. Access to galectin-3 inhibitors from chemoenzymatic synthons. *The Journal of Organic Chemistry* **85**, 16099-16114 (2020).
40 Brun, J., Vasiljevic, S., Gangadharan, B., Hensen, M., Chandran, A. V., Hill, M. L., Kiappes, J., Dwek, R. A., Alonzi, D. S. & Struwe, W. B. Analysis of SARS-CoV-2 spike glycosylation reveals shedding of a vaccine candidate. *bioRxiv*, 2020.2011. 2016.384594 (2020).

41 Cui, J. Y., Zhang, F., Nierzwicki, L., Palermo, G., Linhardt, R. J. & Lisi, G. P. Mapping the structural and dynamic determinants of pH-sensitive heparin binding to granulocyte macrophage colony stimulating factor. *Biochemistry* **59**, 3541-3553 (2020).
42 Prasad Patro, L. P., Sudhakar, K. U. & Rathinavelan, T. K-PAM: A unified platform to distinguish Klebsiella species K-and O-antigen types, model antigen structures and identify hypervirulent strains. *bioRxiv*, 2020.2003. 2021.001370 (2020).
43 Kav, B., Grafmüller, A., Schneck, E. & Weikl, T. R. Weak carbohydrate–carbohydrate interactions in membrane adhesion are fuzzy and generic. *Nanoscale* **12**, 17342-17353 (2020).
44 Patel, M. & Sergeev, Y. Functional in silico analysis of human tyrosinase and OCA1 associated mutations. *Journal of analytical & pharmaceutical research* **9**, 81 (2020).
45 Sýkorová, P., Novotná, J., Demo, G., Pompidor, G., Dubská, E., Komárek, J., Fujdiarová, E., Houser, J., Hároníková, L. & Varrot, A. Characterization of novel lectins from Burkholderia pseudomallei and Chromobacterium violaceum with seven-bladed β-propeller fold. *International journal of biological macromolecules* **152**, 1113-1124 (2020).
46 Machín, B., Chaves, S., Ávila, C., Pera, L. M., Chehín, R. N. & Pingitore, E. V. Highly reusable invertase biocatalyst: Biological fibrils functionalized by photocrosslinking. *Food chemistry* **331**, 127322 (2020).
47 Kirilin, E. & Švedas, V. Analysis of Glycosyl-Enzyme Intermediate Formation in the Catalytic Mechanism of Influenza Virus Neuraminidase Using Molecular Modeling. *Biochemistry (Moscow)* **85**, 490-498 (2020).
48 Callender, J. A., Sevillano, A. M., Soldau, K., Kurt, T. D., Schumann, T., Pizzo, D. P., Altmeppen, H., Glatzel, M., Esko, J. D. & Sigurdson, C. J. Prion protein post-translational modifications modulate heparan sulfate binding and limit aggregate size in prion disease. *Neurobiology of disease* **142**, 104955 (2020).
49 Ishida, T., Parks, J. M. & Smith, J. C. Insight into the catalytic mechanism of GH11 xylanase: computational analysis of substrate distortion based on a neutron structure. *Journal of the American Chemical Society* **142**, 17966-17980 (2020).
50 Bupp, C. R. *A Nano-Hilic-Ms Platform for Separation and Characterization of Glycoproteins*, Purdue University, (2020).
51 Hira, D., Onoue, T. & Oka, T. Structural basis for the core-mannan biosynthesis of cell wall fungal-type galactomannan in Aspergillus fumigatus. *Journal of Biological Chemistry* **295**, 15407-15417 (2020).
52 Trastoy, B., Naegeli, A., Anso, I., Sjögren, J. & Guerin, M. E. Structural basis of mammalian mucin processing by the human gut O-glycopeptidase OgpA from Akkermansia muciniphila. *Nature communications* **11**, 4844 (2020).
53 Phakeenuya, V., Ratanakhanokchai, K., Kosugi, A. & Tachaapaikoon, C. A novel multifunctional GH9 enzyme from Paenibacillus curdlanolyticus B-6 exhibiting endo/exo functions of cellulase, mannanase and xylanase activities. *Applied microbiology and biotechnology* **104**, 2079-2096 (2020).
54 Campbell, I. R., Dong, Z., Grandgeorge, P., Jimenez, A. M., Rhodes, E. R., Lee, E., Edmundson, S., Subban, C. V., Sprenger, K. G. & Roumeli, E. The Role of Biomolecular Building Blocks on the Cohesion of Biomatter Plastics. *Available at SSRN 4734573* (2020).
55 Soliman, C., Chua, J. X., Vankemmelbeke, M., McIntosh, R. S., Guy, A. J., Spendlove, I., Durrant, L. G. & Ramsland, P. A. The terminal sialic acid of stage-specific embryonic antigen-4 has a crucial role in binding to a cancer-targeting antibody. *Journal of Biological Chemistry* **295**, 1009-1020 (2020).
56 Ilmjärv, S., Abdul, F., Acosta-Gutiérrez, S., Estarellas, C., Galdadas, I., Casimir, M., Alessandrini, M., Gervasio, F. L. & Krause, K.-H. Epidemiologically most successful

SARS-CoV-2 variant: concurrent mutations in RNA-dependent RNA polymerase and spike protein. *MedRxiv*, 2020.2008. 2023.20180281 (2020).
57 Atack, J. M., Day, C. J., Poole, J., Brockman, K. L., Timms, J. R., Winter, L. E., Haselhorst, T., Bakaletz, L. O., Barenkamp, S. J. & Jennings, M. P. The nontypeable Haemophilus influenzae major adhesin hia is a dual-function lectin that binds to human-specific respiratory tract sialic acid glycan receptors. *Mbio* **11**, 10.1128/mbio. 02714-02720 (2020).
58 Bernardi, A., Huang, Y., Harris, B., Xiong, Y., Nandi, S., McDonald, K. A. & Faller, R. Development and simulation of fully glycosylated molecular models of ACE2-Fc fusion proteins and their interaction with the SARS-CoV-2 spike protein binding domain. *PLoS One* **15**, e0237295 (2020).
59 Besançon, C., Guillot, A., Blaise, S., Dauchez, M., Belloy, N., Prévoteau-Jonquet, J. & Baud, S. Umbrella Visualization: A method of analysis dedicated to glycan flexibility with UnityMol. *Methods* **173**, 94-104 (2020).
60 Besançon, C., Wong, H., Rao, R., Dauchez, M., Belloy, N., Prévoteau-Jonquet, J. & Baud, S. in *Workshop on Molecular Graphics and Visual Analysis of Molecular Data (MolVA)(2020).*
61 Veluraja, K., Shanmugam, N. R. S., Blessy, J. J., Jeyaram, R. A., Lalithamaheswari, B. & Gromiha, M. M. in *Protein Interactions* 299-332 (2020).
62 Sooyound, L., Yamaguchi, Y. & Sugita, Y. Molecular dynamics simulation of glycans. *Trends Glycosci Glycotechnol* **32**, J93-J98, https://doi.org/10.4052/tigg.1616.1J (2020).
63 Grant, O. C., Montgomery, D., Ito, K. & Woods, R. J. Analysis of the SARS-CoV-2 spike protein glycan shield reveals implications for immune recognition. *Sci Rep* **10**, 14991, https://doi.org/10.1038/s41598-020-71748-7 (2020).
64 Flowers, S. A., Grant, O. C., Woods, R. J. & Rebeck, G. W. O-glycosylation on cerebrospinal fluid and plasma apolipoprotein E differs in the lipid-binding domain. *Glycobiology* **30**, 74-85, https://doi.org/10.1093/glycob/cwz084 (2020).
65 Mandalasi, M., Kim, H. W., Thieker, D., Sheikh, M. O., Gas-Pascual, E., Rahman, K., Zhao, P., Daniel, N. G., van der Wel, H., Ichikawa, H. T., Glushka, J. N., Wells, L., Woods, R. J., Wood, Z. A. & West, C. M. A terminal α3-galactose modification regulates an E3 ubiquitin ligase subunit in Toxoplasma gondii. *J Biol Chem* **295**, 9223-9243, https://doi.org/10.1074/jbc.RA120.013792 (2020).
66 Tang, M., Wang, X., Gandhi, N. S., Foley, B. L., Burrage, K., Woods, R. J. & Gu, Y. Effect of hydroxylysine-O-glycosylation on the structure of type I collagen molecule: A computational study. *Glycobiology* **30**, 830-843, https://doi.org/10.1093/glycob/cwaa026 (2020).
67 Kim, S. Y., Jin, W., Sood, A., Montgomery, D. W., Grant, O. C., Fuster, M. M., Fu, L., Dordick, J. S., Woods, R. J., Zhang, F. & Linhardt, R. J. Characterization of heparin and severe acute respiratory syndrome-related coronavirus 2 (SARS-CoV-2) spike glycoprotein binding interactions. *Antiviral Res* **181**, 104873, https://doi.org/10.1016/j.antiviral.2020.104873 (2020).
68 Zhao, P., Praissman, J. L., Grant, O. C., Cai, Y., Xiao, T., Rosenbalm, K. E., Aoki, K., Kellman, B. P., Bridger, R., Barouch, D. H., Brindley, M. A., Lewis, N. E., Tiemeyer, M., Chen, B., Woods, R. J. & Wells, L. Virus-Receptor Interactions of Glycosylated SARS-CoV-2 Spike and Human ACE2 Receptor. *bioRxiv*, https://doi.org/10.1101/2020.06.25.172403 (2020).
69 Olson, L. J., Misra, S. K., Ishihara, M., Battaile, K. P., Grant, O. C., Sood, A., Woods, R. J., Kim, J. P., Tiemeyer, M., Ren, G., Sharp, J. S. & Dahms, N. M. Allosteric regulation of lysosomal enzyme recognition by the cation-independent mannose 6-phosphate receptor. *Commun Biol* **3**, 498, https://doi.org/10.1038/s42003-020-01211-w (2020).

70 Shao, C., Feng, Z., Westbrook, J. D., Peisach, E., Berrisford, J., Ikegawa, Y., Kurisu, G., Velankar, S., Burley, S. K. & Young, J. Y. Modernized uniform representation of carbohydrate molecules in the Protein Data Bank. *Glycobiology* **31**, 1204-1218 (2021).
71 Fogarty, C. A. & Fadda, E. Oligomannose N-Glycans 3D Architecture and Its Response to the FcγRIIIa Structural Landscape. *The journal of physical chemistry B* **125**, 2607-2616 (2021).
72 Chen, N., Chen, L., He, Q., Sun, Q. & Zeng, W.-C. Effects of tea polyphenols on physicochemical properties of wheat starch and bread quality and their action mechanism. (2021).
73 Novakovic, M., Battistel, M. D., Azurmendi, H. F., Concilio, M.-G., Freedberg, D. n. I. & Frydman, L. The incorporation of labile protons into multidimensional NMR analyses: glycan structures revisited. *Journal of the American Chemical Society* **143**, 8935-8948 (2021).
74 Plazinska, A. & Plazinski, W. Comparison of carbohydrate force fields in molecular dynamics simulations of protein–carbohydrate complexes. *Journal of chemical theory and computation* **17**, 2575-2585 (2021).
75 Reiding, K. R., Lin, Y.-H., van Alphen, F. P., Meijer, A. B. & Heck, A. J. Neutrophil azurophilic granule glycoproteins are distinctively decorated by atypical pauci-and phosphomannose glycans. *Communications Biology* **4**, 1012 (2021).
76 Dhakal, R., Nieman, R., Valente, D. C., Cardozo, T. M., Jayee, B., Aqdas, A., Peng, W., Aquino, A. J., Mechref, Y. & Lischka, H. A general new method for calculating the molecular nonpolar surface for analysis of LC-MS data. *International journal of mass spectrometry* **461**, 116495 (2021).
77 Mohammed, S. & Ferry, N. Characterization of sialic acid affinity of the binding domain of mistletoe lectin isoform one. *International Journal of Molecular Sciences* **22**, 8284 (2021).
78 Sang, S., Xu, X., Zhu, X. & Narsimhan, G. Complexation of 26-Mer amylose with egg yolk lipids with different numbers of tails using a molecular dynamics simulation. *Foods* **10**, 2355 (2021).
79 Kadowaki, M. A., Briganti, L., Evangelista, D. E., Echevarría-Poza, A., Tryfona, T., Pellegrini, V. O., Nakayama, D. G., Dupree, P. & Polikarpov, I. Unlocking the structural features for the xylobiohydrolase activity of an unusual GH11 member identified in a compost-derived consortium. *Biotechnology and Bioengineering* **118**, 4052-4064 (2021).
80 Chakraborty, S., Wagh, K., Gnanakaran, S. & López, C. A. Development of Martini 2.2 parameters for N-glycans: A case study of the HIV-1 Env glycoprotein dynamics. *Glycobiology* **31**, 787-799 (2021).
81 Tagliamonte, M., Abid, N., Borocci, S., Sangiovanni, E. & Ostrov, D. Multiple recombination events and strong purifying selection at the origin of SARS-CoV-2 spike glycoprotein increased correlated dynamic movements. *Int. J. Mol. Sci* **22**, 80 (2021).
82 Lopez Bautista, C. A., Wagh, K., Chakraborty, S. & Gnanakaran, S. Development of Martini 2.2 parameters for N-glycans: a case study of the HIV-1 Env glycoprotein dynamics. *Glycobiology (Online)* (2021).
83 Datta, A. K. & Sukhija, N. in *Glycome: The Hidden Code in Biology. Hauppauge, NY: Nova Science Publishers. p* 323-375 (2021).
84 Brun, J., Vasiljevic, S. a., Gangadharan, B., Hensen, M., V. Chandran, A., Hill, M. L., Kiappes, J., Dwek, R. A., Alonzi, D. S. & Struwe, W. B. Assessing antigen structural integrity through glycosylation analysis of the SARS-CoV-2 viral spike. *ACS central science* **7**, 586-593 (2021).
85 Unione, L., Ardá, A., Jiménez-Barbero, J. & Millet, O. NMR of glycoproteins: profiling, structure, conformation and interactions. *Current Opinion in Structural Biology* **68**, 9-17 (2021).

86 Marchetti, R., Forgione, R. E., Fabregat, F. N., Di Carluccio, C., Molinaro, A. & Silipo, A. Solving the structural puzzle of bacterial glycome. *Current opinion in structural biology* **68**, 74-83 (2021).
87 Dias, J. A. & Ulloa-Aguirre, A. New human follitropin preparations: how glycan structural differences may affect biochemical and biological function and clinical effect. *Frontiers in endocrinology* **12**, 636038 (2021).
88 Lukassen, M. V., Franc, V., Hevler, J. F. & Heck, A. J. Similarities and differences in the structures and proteoform profiles of the complement proteins C6 and C7. *Proteomics* **21**, 2000310 (2021).
89 Broszeit, F., van Beek, R. J., Unione, L., Bestebroer, T. M., Chapla, D., Yang, J.-Y., Moremen, K. W., Herfst, S., Fouchier, R. A. & de Vries, R. P. Glycan remodeled erythrocytes facilitate antigenic characterization of recent A/H3N2 influenza viruses. *Nature Communications* **12**, 5449 (2021).
90 Carneiro, R. F., Aguiar, E. S., Santos, V. F., Santos, A. L., Santos, M. H., Roma, R. R., Silva, R. R., Leal, M. L., Silva, L. T. & Rocha, B. A. Elucidation of the primary structure and molecular modeling of Parkia pendula lectin and in vitro evaluation of the leishmanicidal activity. *Process Biochemistry* **101**, 1-10 (2021).
91 Zdunek, A., Pieczywek, P. M. & Cybulska, J. The primary, secondary, and structures of higher levels of pectin polysaccharides. *Compr Rev Food Sci F* **20**, 1101-1117 (2021).
92 Marques, G. F. O., Pires, A. F., Osterne, V. J. S., Pinto-Junior, V. R., Silva, I. B., Martins, M. G. Q., Oliveira, M. V., Gomes, A. M., de Souza, L. A. G. & Pavão, M. S. G. Vatairea guianensis lectin stimulates changes in gene expression and release of TNF-α from rat peritoneal macrophages via glycoconjugate binding. *Journal of Molecular Recognition* **34**, e2922 (2021).
93 Nguyen, T. B., Lane, D. P. & Verma, C. S. Can glycosylation mask the detection of MHC expressing p53 peptides by T cell receptors? *Biomolecules* **11**, 1056 (2021).
94 Bupp, C. R., Schwartz, C., Wei, B. & Wirth, M. J. Protein-induced conformational change in glycans decreases the resolution of glycoproteins in hydrophilic interaction liquid chromatography. *Journal of separation science* **44**, 1581-1591 (2021).
95 Mir, S., Ashraf, S., Saeed, M., Rahman, A.-u. & Ul-Haq, Z. Protonation states at different pH, conformational changes and impact of glycosylation in synapsin Ia. *Physical Chemistry Chemical Physics* **23**, 16718-16729 (2021).
96 Kotani, O., Suzuki, Y., Saito, S., Ainai, A., Ueno, A., Hemmi, T., Sano, K., Tabata, K., Yokoyama, M. & Suzuki, T. Structure-Guided Creation of an Anti-HA Stalk Antibody F11 Derivative That Neutralizes Both F11-Sensitive and-Resistant Influenza A (H1N1) pdm09 Viruses. *Viruses* **13**, 1733 (2021).
97 Botelho, T., Osterne, V. J., Pinto-Junior, V. R., Oliveira, M. V., Cavada, B. S., Nascimento, K. S. & Dos Santos, L. Differential vasodilator effect of Dioclea rostrata lectin in conductance and resistance arteries: Mechanisms and glycoconjugate binding relationships. *Basic & Clinical Pharmacology & Toxicology* **129**, 130-138 (2021).
98 Gerwig, G. J. & Gerwig, G. J. in *The Art of Carbohydrate Analysis Techniques in Life Science and Biomedicine for the Non-Expert (TLSBNE)* Ch. Glycobioinformatics, 297-312 (2021).
99 Wu, J., Tan, Z., Li, H., Lin, M., Jiang, Y., Liang, L., Ma, Q., Gou, J., Ning, L. & Li, X. Melatonin reduces proliferation and promotes apoptosis of bladder cancer cells by suppressing O-GlcNAcylation of cyclin-dependent-like kinase 5. *Journal of Pineal Research* **71**, e12765 (2021).
100 Sun, Z., Ren, K., Zhang, X., Chen, J., Jiang, Z., Jiang, J., Ji, F., Ouyang, X. & Li, L. Mass spectrometry analysis of newly emerging coronavirus HCoV-19 spike protein and human ACE2 reveals camouflaging glycans and unique post-translational modifications. *Engineering* **7**, 1441-1451 (2021).

101 Porciúncula-González, C., Cagnoni, A. J., Fontana, C., Mariño, K. V., Saenz-Méndez, P., Giacomini, C. & Irazoqui, G. Structural insights in galectin-1-glycan recognition: Relevance of the glycosidic linkage and the N-acetylation pattern of sugar moieties. *Bioorganic & Medicinal Chemistry* **44**, 116309 (2021).
102 Vessella, G., Marchetti, R., Del Prete, A., Traboni, S., Iadonisi, A., Schiraldi, C., Silipo, A. & Bedini, E. Semisynthetic isomers of fucosylated chondroitin sulfate polysaccharides with fucosyl branches at a non-natural site. *Biomacromolecules* **22**, 5151-5161 (2021).
103 Dwivedi, R., Samanta, P., Sharma, P., Zhang, F., Mishra, S. K., Kucheryavy, P., Kim, S. B., Aderibigbe, A. O., Linhardt, R. J. & Tandon, R. Structural and kinetic analyses of holothurian sulfated glycans suggest potential treatment for SARS-CoV-2 infection. *Journal of Biological Chemistry* **297** (2021).
104 Schuurs, Z. P., Hammond, E., Elli, S., Rudd, T. R., Mycroft-West, C. J., Lima, M. A., Skidmore, M. A., Karlsson, R., Chen, Y.-H. & Bagdonaite, I. Evidence of a putative glycosaminoglycan binding site on the glycosylated SARS-CoV-2 spike protein N-terminal domain. *Computational and Structural Biotechnology Journal* **19**, 2806-2818 (2021).
105 Miller, N. L., Clark, T., Raman, R. & Sasisekharan, R. Glycans in virus-host interactions: A structural perspective. *Frontiers in molecular biosciences* **8**, 666756 (2021).
106 Paiardi, G., Milanesi, M., Wade, R. C., D'Ursi, P. & Rusnati, M. A Bittersweet Computational journey among glycosaminoglycans. *Biomolecules* **11**, 739 (2021).
107 Zhang, S., Li, C., Gilbert, R. G. & Malde, A. K. Understanding the binding of starch fragments to granule-bound starch synthase. *Biomacromolecules* **22**, 4730-4737 (2021).
108 Li, Y., Wang, T., Zhang, J., Shao, B., Gong, H., Wang, Y., He, X., Liu, S. & Liu, T. Y. Exploring the Regulatory Function of the N-terminal Domain of SARS-CoV-2 Spike Protein through Molecular Dynamics Simulation. *Advanced theory and simulations* **4**, 2100152 (2021).
109 Gao, M., Li, H., Ye, C., Chen, K., Jiang, H. & Yu, K. Glycan epitopes and potential glycoside antagonists of DC-SIGN Involved in COVID-19: in silico study. *Biomolecules* **11**, 1586 (2021).
110 Quintana, J. I., Delgado, S., Núñez-Franco, R., Cañada, F. J., Jiménez-Osés, G., Jiménez-Barbero, J. & Ardá, A. Galectin-4 N-terminal domain: binding preferences toward a and B antigens with different peripheral core presentations. *Frontiers in chemistry* **9**, 664097 (2021).
111 Perez, S., Fadda, E. & Makshakova, O. Computational modeling in glycoscience. *Comprehensive Glycoscience: Second Edition*, 374-404 (2021).
112 Motamedi, Z., Rajabi-Maham, H. & Azimzadeh Irani, M. Glycosylation promotes the cancer regulator EGFR-ErbB2 heterodimer formation—molecular dynamics study. *Journal of Molecular Modeling* **27**, 1-14 (2021).
113 García-García, A., Serna, S., Yang, Z., Delso, I., Taleb, V., Hicks, T., Artschwager, R., Vakhrushev, S. Y., Clausen, H. & Angulo, J. FUT8-directed core fucosylation of N-glycans is regulated by the glycan structure and protein environment. *ACS catalysis* **11**, 9052-9065 (2021).
114 Soares, C. O., Grosso, A. S., Ereño-Orbea, J., Coelho, H. & Marcelo, F. Molecular recognition insights of sialic acid glycans by distinct receptors unveiled by NMR and molecular modeling. *Frontiers in Molecular Biosciences* **8**, 727847 (2021).
115 Cabezas-Pérusse, Y., Daligault, F., Ferrières, V., Tasseau, O. & Tranchimand, S. Modulation of the Activity and Regioselectivity of a Glycosidase: Development of a Convenient Tool for the Synthesis of Specific Disaccharides. *Molecules* **26**, 5445 (2021).
116 Wu, Q., Zhang, C., Zhang, K., Chen, Q., Wu, S., Huang, H., Huang, T., Zhang, N., Wang, X. & Li, W. ppGalNAc-T4-catalyzed O-Glycosylation of TGF-β type Ⅱ receptor regulates breast cancer cells metastasis potential. *JBC* **296** (2021).

117 Prestegard, J. H. A perspective on the PDB's impact on the field of glycobiology. *Journal of Biological Chemistry* **296** (2021).
118 Paiardi, G., Richter, S., Oreste, P., Urbinati, C., Rusnati, M. & Wade, R. C. Three-fold mechanism of inhibition of SARS-CoV-2 infection by the interaction of the spike glycoprotein with heparin. *arXiv preprint arXiv:2103.07722* (2021).
119 Fadda, E. Understanding the structure and function of viral glycosylation by molecular simulations: state-of-the-art and recent case studies. *Comprehensive Glycoscience*, 405 (2021).
120 Sundar, S., Sandilya, A. A. & Priya, M. H. Unraveling the influence of osmolytes on water hydrogen-bond network: From local structure to graph theory analysis. *Journal of Chemical Information and Modeling* **61**, 3927-3944 (2021).
121 Camacho, E., Dong, Y., Anglero-Rodriguez, Y., Smith, D., de Souza Jacomini, R., Dimopoulos, G. & Casadevall, A. Analysis of melanotic Plasmodium spp. capsules in mosquitoes reveal eumelanin-pheomelanin composition and identify Ag Mesh as a modulator of parasite infection. *bioRxiv*, 2021.2005. 2007.443077 (2021).
122 Knoppova, B., Reily, C., King, R. G., Julian, B. A., Novak, J. & Green, T. J. Pathogenesis of IgA nephropathy: current understanding and implications for development of disease-specific treatment. *Journal of Clinical Medicine* **10**, 4501 (2021).
123 Jayaprakash, N. G. & Surolia, A. Spike protein and the various cell-surface carbohydrates: an interaction study. *ACS chemical biology* **17**, 103-117 (2021).
124 Paiardi, G. *Glycan-Protein interactions in pathological processes: development of new molecular and cellular models for glycobiology-oriented studies* PhD thesis, (2021).
125 Rasin, A. B., Shevchenko, N. M., Silchenko, A. S., Kusaykin, M. I., Likhatskaya, G. N., Zvyagintseva, T. N. & Ermakova, S. P. Relationship between the structure of a highly regular fucoidan from Fucus evanescens and its ability to form nanoparticles. *International Journal of Biological Macromolecules* **185**, 679-687 (2021).
126 Trastoy, B., Du, J. J., Li, C., García-Alija, M., Klontz, E. H., Roberts, B. R., Donahue, T. C., Wang, L.-X., Sundberg, E. J. & Guerin, M. E. GH18 endo-β-N-acetylglucosaminidases use distinct mechanisms to process hybrid-type N-linked glycans. *Journal of Biological Chemistry* **297** (2021).
127 Rajendran, M., Ferran, M. C. & Babbitt, G. A. Identifying protein sites contributing to vaccine escape via statistical comparisons of short-term molecular dynamics simulations. *bioRxiv*, 2021.2012. 2006.471374 (2021).
128 Giron, C. C., Laaksonen, A. & Barroso da Silva, F. L. Up state of the SARS-COV-2 spike homotrimer favors an increased virulence for new variants. *Frontiers in medical technology* **3**, 694347 (2021).
129 Martí-Marí, O., Martínez-Gualda, B., de la Puente-Secades, S., Mills, A., Quesada, E., Abdelnabi, R., Sun, L., Boonen, A., Noppen, S. & Neyts, J. Double arylation of the indole side chain of tri-and tetrapodal tryptophan derivatives renders highly potent HIV-1 and EV-A71 entry inhibitors. *Journal of Medicinal Chemistry* **64**, 10027-10046 (2021).
130 Sreenivasan, C. C., Sheng, Z., Wang, D. & Li, F. Host Range, Biology, and Species Specificity of Seven-Segmented Influenza Viruses—A Comparative Review on Influenza C and D. *Pathogens* **10**, 1583 (2021).
131 Ilmjärv, S., Abdul, F., Acosta-Gutiérrez, S., Estarellas, C., Galdadas, I., Casimir, M., Alessandrini, M., Gervasio, F. L. & Krause, K.-H. Concurrent mutations in RNA-dependent RNA polymerase and spike protein emerged as the epidemiologically most successful SARS-CoV-2 variant. *Scientific reports* **11**, 13705 (2021).
132 Fernández de Toro Ronda, B. *New methologies for studying carbohydrate-protein interacions by nuclear magnetic resonance* PhD thesis, (2021).
133 Hosoda, M. & Kinoshita, S. An introduction to glycan-related informatics. *JSBi Bioinformatics Review* **2**, 87-95 (2021).

134 Nascimento, T. B. Vasodilation mediated by Dioclea lectin rostrata in conductance arteries and resistance: mechanisms and predictions of binding with glycoconjugates. (2021).
135 Abaramak, G., Porras-Domínguez, J. R., Janse van Rensburg, H. C., Lescrinier, E., Toksoy Öner, E., Kırtel, O. & Van den Ende, W. Functional and molecular characterization of the Halomicrobium sp. IBSBa Inulosucrase. *Microorganisms* **9**, 749 (2021).
136 Akinbiyi, E. O., Abramowitz, L. K., Bauer, B. L., Stoll, M. S., Hoppel, C. L., Hsiao, C.-P., Hanover, J. A. & Mears, J. A. Blocked O-GlcNAc cycling alters mitochondrial morphology, function, and mass. *Scientific Reports* **11**, 22106 (2021).
137 Basu, S., Chakravarty, D., Bhattacharyya, D., Saha, P. & Patra, H. K. Plausible blockers of Spike RBD in SARS-CoV2—Molecular design and underlying interaction dynamics from high-level structural descriptors. *Journal of Molecular Modeling* **27**, 191 (2021).
138 Borocci, S., Cerchia, C., Grottesi, A., Sanna, N., Prandi, I. G., Abid, N., Beccari, A. R., Chillemi, G. & Talarico, C. Altered local interactions and long-range communications in UK variant (B. 1.1. 7) spike glycoprotein. *International Journal of Molecular Sciences* **22**, 5464 (2021).
139 Breslawec, A. P., Wang, S., Li, C. & Poulin, M. B. Anionic amino acids support hydrolysis of poly-β-(1, 6)-N-acetylglucosamine exopolysaccharides by the biofilm dispersing glycosidase Dispersin B. *Journal of Biological Chemistry* **296** (2021).
140 Tjondro, H. C., Ugonotti, J., Kawahara, R., Chatterjee, S., Loke, I., Chen, S., Soltermann, F., Hinneburg, H., Parker, B. L., Venkatakrishnan, V., Dieckmann, R., Grant, O. C., Bylund, J., Rodger, A., Woods, R. J., Karlsson-Bengtsson, A., Struwe, W. B. & Thaysen-Andersen, M. Hyper-truncated Asn355- and Asn391-glycans modulate the activity of neutrophil granule myeloperoxidase. *J Biol Chem* **296**, 100144, https://doi.org/10.1074/jbc.RA120.016342 (2021).
141 French, A. D., Montgomery, D. W., Prevost, N. T., Edwards, J. V. & Woods, R. J. Comparison of cellooligosaccharide conformations in complexes with proteins with energy maps for cellobiose. *Carbohydr Polym* **264**, 118004, https://doi.org/10.1016/j.carbpol.2021.118004 (2021).
142 Fernandez, F. J., Santos-Lopez, J., Martinez-Barricarte, R., Querol-Garcia, J., Martin-Merinero, H., Navas-Yuste, S., Savko, M., Shepard, W. E., Rodriguez de Cordoba, S. & Vega, M. C. The crystal structure of iC3b-CR3 alphaI reveals a modular recognition of the main opsonin iC3b by the CR3 integrin receptor. *Nat Commun* **13**, 1955, https://doi.org/10.1038/s41467-022-29580-2 (2022).
143 Roy, R., Poddar, S. & Kar, P. Comparison of the conformational dynamics of an N-glycan in implicit and explicit solvents. *Carbohydrate Research* **522**, 108700 (2022).
144 Nagarajan, B. & Desai, U. in *Glycosaminoglycans: Methods and Protocols* 49-62 (2022).
145 Vanacore, A., Forgione, M. C., Cavasso, D., Nguyen, H. N. A., Molinaro, A., Saenz, J. P., D'Errico, G., Paduano, L., Marchetti, R. & Silipo, A. Role of EPS in mitigation of plant abiotic stress: The case of Methylobacterium extorquens PA1. *Carbohydrate Polymers* **295**, 119863 (2022).
146 Romeo, I., Prandi, I. G., Giombini, E., Gruber, C. E. M., Pietrucci, D., Borocci, S., Abid, N., Fava, A., Beccari, A. R. & Chillemi, G. The Spike Mutants website: a worldwide used resource against SARS-CoV-2. *International Journal of Molecular Sciences* **23**, 13082 (2022).
147 Wu, L., Peng, C., Yang, Y., Shi, Y., Zhou, L., Xu, Z. & Zhu, W. Exploring the immune evasion of SARS-CoV-2 variant harboring E484K by molecular dynamics simulations. *Brief. Bioinform.* **23**, bbab383 (2022).

148 Meredith, R. J., Tetrault, T., Yoon, M.-K., Zhang, W., Carmichael, I. & Serianni, A. S. N-Acetyl side-chain conformation in saccharides: Solution models obtained from MA'AT analysis. *The Journal of Organic Chemistry* **87**, 8368-8379 (2022).
149 Pirone, L., Nieto-Fabregat, F., Di Gaetano, S., Capasso, D., Russo, R., Traboni, S., Molinaro, A., Iadonisi, A., Saviano, M. & Marchetti, R. Exploring the Molecular Interactions of Symmetrical and Unsymmetrical Selenoglycosides with Human Galectin-1 and Galectin-3. *International Journal of Molecular Sciences* **23**, 8273 (2022).
150 Tetrault, T., Meredith, R. J., Zhang, W., Carmichael, I. & Serianni, A. S. One-Bond 13C–1H and 13C–13C Spin-Coupling Constants as Constraints in MA'AT Analysis of Saccharide Conformation. *The Journal of Physical Chemistry B* **126**, 9506-9515 (2022).
151 Kong, Y., Li, L. & Fu, S. Insights from molecular dynamics simulations for interaction between cellulose microfibrils and hemicellulose. *Journal of Materials Chemistry A* **10**, 14451-14459 (2022).
152 Chen, N., Gao, H.-X., He, Q. & Zeng, W.-C. Wheat starch modified with Ligustrum robustum (Rxob.) Blume extract and its action mechanism. *Foods* **11**, 3187 (2022).
153 Meredith, R. J., McGurn, M., Euell, C., Rutkowski, P., Cook, E., Carmichael, I. & Serianni, A. S. MA'AT Analysis of Aldofuranosyl Rings: Unbiased Modeling of Conformational Equilibria and Dynamics in Solution. *Biochemistry* **61**, 239-251 (2022).
154 Williams, R. V., Huang, C., McDermott, C., Ahmed, T., Columbus, L., Moremen, K. W., Prestegard, J. H. & Amster, I. J. Site-to-site cross-talk in OST-B glycosylation of hCEACAM1-IgV. *Proceedings of the National Academy of Sciences* **119**, e2202992119 (2022).
155 Huang, H., Hou, X., Xu, R., Deng, Z., Wang, Y., Du, G., Rao, Y., Chen, J. & Kang, Z. Structure and cleavage pattern of a hyaluronate 3-glycanohydrolase in the glycoside hydrolase 79 family. *Carbohydrate Polymers* (2022).
156 Versluys, M., Porras-Domínguez, J. R., De Coninck, T., Van Damme, E. J. & Van den Ende, W. A novel chicory fructanase can degrade common microbial fructan product profiles and displays positive cooperativity. *Journal of Experimental Botany* **73**, 1602-1622 (2022).
157 Prandi, I. G., Mavian, C., Giombini, E., Gruber, C. E., Pietrucci, D., Borocci, S., Abid, N., Beccari, A. R., Talarico, C. & Chillemi, G. Structural Evolution of Delta (B. 1.617. 2) and Omicron (BA. 1) Spike Glycoproteins. *International Journal of Molecular Sciences* **23**, 8680 (2022).
158 Purushotham, P., Ho, R., Yu, L., Fincher, G. B., Bulone, V. & Zimmer, J. Mechanism of mixed-linkage glucan biosynthesis by barley cellulose synthase–like CslF6 (1, 3; 1, 4)-β-glucan synthase. *Science Advances* **8**, eadd1596 (2022).
159 Hlima, H. B., Farhat, A., Akermi, S., Khemakhem, B., Halima, Y. B., Michaud, P., Fendri, I. & Abdelkafi, S. In silico evidence of antiviral activity against SARS-CoV-2 main protease of oligosaccharides from Porphyridium sp. *Science of the Total Environment* **836**, 155580 (2022).
160 Kim, H. W. *Structural Insights into How Glycosylation Controls Skp1 in Protists* PhD thesis, University of Georgia, (2022).
161 Roy, S., Ghosh, P., Bandyopadhyay, A. & Basu, S. Capturing a crucial 'disorder-to-order transition'at the heart of the coronavirus molecular pathology—triggered by highly persistent, interchangeable salt-bridges. *Vaccines* **10**, 301 (2022).
162 Lian, D., Zhuang, S., Shui, C., Zheng, S., Ma, Y., Sun, Z., Porras-Domínguez, J. R., Öner, E. T., Liang, M. & Van den Ende, W. Characterization of inulolytic enzymes from the Jerusalem artichoke–derived Glutamicibacter mishrai NJAU-1. *Applied Microbiology and Biotechnology* **106**, 5525-5538 (2022).

163 Xu, L., Zhao, Z.-X., Huang, Y.-A. & Zhu, Q.-J. Preparation of chitosan molecularly imprinted polymers and the recognition mechanism for adsorption of alpha-lipoic acid. *Molecules* **25**, 312 (2022).
164 Uslupehlivan, M. & Uslupehlivan, E. Ş. Glycoinformatics approach for identifying target positions to inhibit initial binding of SARS-CoV-2 S1 protein to the host cell. *Journal of Applied Biological Sciences* **16**, 89-101 (2022).
165 Holmes, S. G., Nagarajan, B. & Desai, U. R. 3-O-Sulfation induces sequence-specific compact topologies in heparan sulfate that encode a dynamic sulfation code. *Computational and Structural Biotechnology Journal* (2022).
166 Pang, Y. T., Acharya, A., Lynch, D. L., Pavlova, A. & Gumbart, J. C. SARS-CoV-2 spike opening dynamics and energetics reveal the individual roles of glycans and their collective impact. *Communications Biology* **5**, 1170 (2022).
167 Lazar, R. D., Akher, F. B., Ravenscroft, N. & Kuttel, M. M. Carbohydrate force fields: the role of small partial atomic charges in preventing conformational collapse. *Journal of Chemical Theory and Computation* **18**, 1156-1172 (2022).
168 Roe, D. R. & Bergonzo, C. prepareforleap: An automated tool for fast PDB-to-parameter generation. *Journal of Computational Chemistry* **43**, 930-935 (2022).
169 Terrasan, C. R. F., Rubio, M. V., Gerhardt, J. A., Cairo, J. P. F., Contesini, F. J., Zubieta, M. P., Figueiredo, F. L. d., Valadares, F. L., Corrêa, T. L. R. & Murakami, M. T. Deletion of AA9 lytic polysaccharide monooxygenases impacts A. nidulans secretome and growth on lignocellulose. *Microbiology spectrum* **10**, e02125-02121 (2022).
170 Li, Z., Liu, L., Unione, L., Lang, Y., de Groot, R. J. & Boons, G.-J. Synthetic O-acetyl-N-glycolylneuraminic acid oligosaccharides reveal host-associated binding patterns of coronaviral glycoproteins. *ACS Infectious Diseases* **8**, 1041-1050 (2022).
171 Peng, W., Rayaprolu, V., Parvate, A. D., Pronker, M. F., Hui, S., Parekh, D., Shaffer, K., Yu, X., Saphire, E. O. & Snijder, J. Glycan shield of the ebolavirus envelope glycoprotein GP. *Communications biology* **5**, 785 (2022).
172 Butnev, V. Y., May, J. V., Brown, A. R., Sharma, T., Butnev, V. Y., White, W. K., Harvey, D. J. & Bousfield, G. R. Human FSH Glycoform α-Subunit Asparagine52 Glycans: Major Glycan Structural Consistency, Minor Glycan Variation in Abundance. *Frontiers in Endocrinology* **13**, 767661 (2022).
173 Mariethoz, J., Alocci, D., Karlsson, N. G., Packer, N. H. & Lisacek, F. An Interactive View of Glycosylation. *Glycosylation: Methods and Protocols*, 41-65 (2022).
174 Stagnoli, S., Peccati, F., Connell, S. R., Martinez-Castillo, A., Charro, D., Millet, O., Bruzzone, C., Palazon, A., Ardá, A. & Jiménez-Barbero, J. Assessing the mobility of severe acute respiratory syndrome Coronavirus-2 spike protein glycans by structural and computational methods. *Frontiers in microbiology* **13**, 870938 (2022).
175 Fu, Y., Ning, L., Feng, J., Yu, X., Guan, F. & Li, X. Dynamic regulation of O-GlcNAcylation and phosphorylation on STAT3 under hypoxia-induced EMT. *Cellular Signalling* **93**, 110277 (2022).
176 Izumida, M., Kotani, O., Hayashi, H., Smith, C., Fukuda, T., Suga, K., Iwao, M., Ishibashi, F., Sato, H. & Kubo, Y. Unique mode of antiviral action of a marine alkaloid against ebola virus and SARS-CoV-2. *Viruses* **14**, 816 (2022).
177 Grünewald, F., Punt, M. H., Jefferys, E. E., Vainikka, P. A., König, M., Virtanen, V., Meyer, T. A., Pezeshkian, W., Gormley, A. J. & Karonen, M. Martini 3 coarse-grained force field for carbohydrates. *Journal of Chemical Theory and Computation* **18**, 7555-7569 (2022).
178 Canales, A., Sastre, J., Orduña, J. M., Pérez-Castells, J., Domínguez, G., van der Woude, R., Nycholat, C. M., Paulson, J. C., Boons, G.-J. & Jiménez-Barbero, J. Revealing the evolution towards complex N-glycan specificities of human H1 influenza A viruses. (2022).

179 Hansen, D. K., Hansen, A. L., Koivisto, J. M., Shuoker, B., Abou Hachem, M., Winther, J. R. & Willemoës, M. Engineering Bifidobacterium longum Endo-α-N-acetylgalactosaminidase for Neu5Acα2-3Galβ1-3GalNAc reactivity on Fetuin. *Archives of Biochemistry and Biophysics* **725**, 109280 (2022).
180 Nagae, M., Hirata, T., Tateno, H., Mishra, S. K., Manabe, N., Osada, N., Tokoro, Y., Yamaguchi, Y., Doerksen, R. J. & Shimizu, T. Discovery of a lectin domain that regulates enzyme activity in mouse N-acetylglucosaminyltransferase-IVa (MGAT4A). *Communications Biology* **5**, 695 (2022).
181 Rajendran, M., Ferran, M. C. & Babbitt, G. A. Identifying vaccine escape sites via statistical comparisons of short-term molecular dynamics. *Biophysical Reports* **2** (2022).
182 Oganesyan, I., Hajduk, J., Harrison, J. A., Marchand, A., Czar, M. F. & Zenobi, R. Exploring gas-phase MS methodologies for structural elucidation of branched N-glycan isomers. *Analytical Chemistry* **94**, 10531-10539 (2022).
183 Perez, S. & Makshakova, O. Multifaceted computational modeling in glycoscience. *Chemical Reviews* **122**, 15914-15970 (2022).
184 Gabrielli, V., Baretta, R., Pilot, R., Ferrarini, A. & Frasconi, M. Insights into the gelation mechanism of metal-coordinated hydrogels by paramagnetic NMR spectroscopy and molecular dynamics. *Macromolecules* **55**, 450-461 (2022).
185 Paiardi, G., Richter, S., Oreste, P., Urbinati, C., Rusnati, M. & Wade, R. C. The binding of heparin to spike glycoprotein inhibits SARS-CoV-2 infection by three mechanisms. *Journal of Biological Chemistry* **298** (2022).
186 Scherbinina, S. I., Frank, M. & Toukach, P. V. Carbohydrate Structure Database oligosaccharide conformation tool. *Glycobiology* **32**, 460-468 (2022).
187 Yu, F., Teng, Y., Yang, S., He, Y., Zhang, Z., Yang, H., Ding, C.-F. & Zhou, P. The thermodynamic and kinetic mechanisms of a Ganoderma lucidum proteoglycan inhibiting hIAPP amyloidosis. *Biophysical Chemistry* **280**, 106702 (2022).
188 Haji, S., Ito, T., Guenther, C., Nakano, M., Shimizu, T., Mori, D., Chiba, Y., Tanaka, M., Mishra, S. K. & Willment, J. A. Human Dectin-1 is O-glycosylated and serves as a ligand for C-type lectin receptor CLEC-2. *Elife* **11**, e83037 (2022).
189 Zhang, C., Kim, E., Cui, J., Wang, Y., Lee, Y. & Zhang, G. Influence of the ecological environment on the structural characteristics and bioactivities of polysaccharides from alfalfa (Medicago sativa L.). *Food Funct* **13**, 7029-7045 (2022).
190 Shofolawe-Bakare, O. T., de Mel, J. U., Mishra, S. K., Hossain, M., Hamadani, C. M., Pride, M. C., Dasanayake, G. S., Monroe, W., Roth, E. W. & Tanner, E. E. ROS-Responsive Glycopolymeric Nanoparticles for Enhanced Drug Delivery to Macrophages. *Macromolecular bioscience* **22**, 2200281 (2022).
191 Stratilová, B., Stratilová, E., Hrmova, M. & Kozmon, S. Definition of the acceptor substrate binding specificity in plant xyloglucan endotransglycosylases using computational chemistry. *International Journal of Molecular Sciences* **23**, 11838 (2022).
192 Chang, T.-H., Gloria, Y. C., Hellmann, M. J., Greve, C. L., Le Roy, D., Roger, T., Bork, F., Bugl, S., Jakob, J. & Kasper, L. Transkingdom mechanism of MAMP generation by chitotriosidase (CHIT1) feeds oligomeric chitin from fungal pathogens and allergens into TLR2-mediated innate immune sensing. *bioRxiv*, 2022.2002. 2017.479713 (2022).
193 Carbajo, D., Pérez, Y., Guerra-Rebollo, M., Prats, E., Bujons, J. & Alfonso, I. Dynamic Combinatorial Optimization of In Vitro and In Vivo Heparin Antidotes. *Journal of medicinal chemistry* **65**, 4865-4877 (2022).
194 Xu, H., Palpant, T., Weinberger, C. & Shaw, D. E. Characterizing receptor flexibility to predict mutations that lead to human adaptation of influenza hemagglutinin. *J. Chem. Theory Comput.* **18**, 4995-5005 (2022).

195 Kadooka, C., Hira, D., Tanaka, Y., Chihara, Y., Goto, M. & Oka, T. Mnt1, an α-(1→ 2)-mannosyltransferase responsible for the elongation of N-glycans and O-glycans in Aspergillus fumigatus. *Glycobiology* **32**, 1137-1152 (2022).
196 Motamedi, Z., Shahsavari, M., Rajabi-Maham, H. & Azimzadeh Irani, M. Cancer regulator EGFR-ErbB4 heterodimer is stabilized through glycans at the dimeric interface. *Journal of Molecular Modeling* **28**, 399 (2022).
197 Masoomi Nomandan, S. Z., Azimzadeh Irani, M. & Hosseini, S. M. In silico design of refined ferritin-SARS-CoV-2 glyco-RBD nanoparticle vaccine. *Frontiers in Molecular Biosciences* **9**, 976490 (2022).
198 García-Alija, M., Du, J. J., Ordóñez, I., Diz-Vallenilla, A., Moraleda-Montoya, A., Sultana, N., Huynh, C. G., Li, C., Donahue, T. C. & Wang, L.-X. Mechanism of cooperative N-glycan processing by the multi-modular endoglycosidase EndoE. *Nature Communications* **13**, 1137 (2022).
199 Koval'ová, T., Kovaľ, T., Stránský, J., Kolenko, P., Dušková, J., Švecová, L., Vodičková, P., Spiwok, V., Benešová, E. & Lipovová, P. The first structure–function study of GH151 α-l-fucosidase uncovers new oligomerization pattern, active site complementation, and selective substrate specificity. *The FEBS Journal* **289**, 4998-5020 (2022).
200 Lalithamaheswari, B. & Anu Radha, C. Structural and conformational dynamics of human milk oligosaccharides, lacto-N-fucopentaose I and II, through molecular dynamics simulation. *Journal of Carbohydrate Chemistry* **41**, 385-404 (2022).
201 Zhou, L., Liu, T., Mo, M., Shi, Y., Wu, L., Li, Y., Qin, Q., Zhu, W., Wu, C. & Gong, L. Exploring the binding affinity and mechanism between ACE2 and the trimers of Delta and Omicron spike proteins by molecular dynamics simulation and bioassay. *J Chem Inf Model* **62**, 4512-4522 (2022).
202 Rajendran, M. & Babbitt, G. A. Persistent cross-species SARS-CoV-2 variant infectivity predicted via comparative molecular dynamics simulation. *Royal Society Open Science* **9**, 220600 (2022).
203 Cui, J., Wang, Y., Kim, E., Zhang, C., Zhang, G. & Lee, Y. Structural characteristics and immunomodulatory effects of a long-chain polysaccharide from Laminaria japonica. *Frontiers in nutrition* **9**, 762595 (2022).
204 Weigle, A. T., Feng, J. & Shukla, D. Thirty years of molecular dynamics simulations on posttranslational modifications of proteins. *Physical Chemistry Chemical Physics* **24**, 26371-26397 (2022).
205 Weyer, R., Hellmann, M. J., Hamer-Timmermann, S. N., Singh, R. & Moerschbacher, B. M. Customized chitooligosaccharide production—controlling their length via engineering of rhizobial chitin synthases and the choice of expression system. *Frontiers in Bioengineering and Biotechnology* **10**, 1073447 (2022).
206 Cao, Y., Qiao, Y., Cui, S. & Ge, J. Origin of metal cluster tuning enzyme activity at the Bio-Nano interface. *JACS Au* **2**, 961-971 (2022).
207 Mukherjee, R., Somovilla, V. J., Chiodo, F., Bruijns, S., Pieters, R. J., Garssen, J., van Kooyk, Y., Kraneveld, A. D. & van Bergenhenegouwen, J. Human Milk Oligosaccharide 2′-Fucosyllactose Inhibits Ligand Binding to C-Type Lectin DC-SIGN but Not to Langerin. *International journal of molecular sciences* **23**, 14745 (2022).
208 Huang, Y. *Atomistic Modeling and Computational Study of Reactive Systems and Biological Applications* PhD thesis, University of California, Davis, (2022).
209 Kappes, E. C. *Molecular Characterization of the Inhibin A Heterodimer and its Function as an Activin Antagonist*, University of Cincinnati, (2022).
210 Mishra, S., Samanta, P. & Doerksen, R. J. *Computational Chemistry and Bioinformatics Research CORE (CCBRC)*, (2022).
211 Harris, B. S. *Molecular Modeling for 3D Printing and Biological Applications* PhD thesis, University of California, Davis, (2022).

212 Capurro, J. I. B. *Study of carbohydrate-binding proteins of biological relevance* PhD thesis, Universidad de Buenos Aires, (2022).
213 Akinbiyi, E. O. *Understanding How O-GlcNAcylation and Phosphorylation Regulates the Mitochondrial Fission Machinery in Glioblastoma* PhD thesis, Case Western Reserve University, (2022).
214 Al Kafri, N., Ahnström, J., Teraz-Orosz, A., Chaput, L., Singh, N., Villoutreix, B. O. & Hafizi, S. The first laminin G-like domain of protein S is essential for binding and activation of Tyro3 receptor and intracellular signalling. *Biochemistry and biophysics reports* **30**, 101263 (2022).
215 Amos, R. A., Atmodjo, M. A., Huang, C., Gao, Z., Venkat, A., Taujale, R., Kannan, N., Moremen, K. W. & Mohnen, D. Polymerization of the backbone of the pectic polysaccharide rhamnogalacturonan I. *Nature plants* **8**, 1289-1303 (2022).
216 Anderson, K. W., Bergonzo, C., Scott, K., Karageorgos, I. L., Gallagher, E. S., Tayi, V. S., Butler, M. & Hudgens, J. W. HDX-MS and MD simulations provide evidence for stabilization of the IgG1—FcγRIa (CD64a) immune complex through intermolecular glycoprotein bonds. *Journal of molecular biology* **434**, 167391 (2022).
217 Aoki-Kinoshita, K. F., Campbell, M. P., Lisacek, F., Neelamegham, S., York, W. S. & Packer, N. H. Glycoinformatics. *Essentials of Glycobiology [Internet]. 4th edition* (2022).
218 Azari-Anpar, M., Degraeve, P., Oulahal, N., Adt, I., Jahanbin, K., Demarigny, Y., Assifaoui, A. & Yazdi, F. T. Interaction of Escherichia coli heat-labile enterotoxin B-pentamer with exopolysaccharides from Leuconostoc mesenteroides P35: Insights from surface plasmon resonance and molecular docking studies. *Food Bioscience* **50**, 102058 (2022).
219 Azimzadeh Irani, M. & Ejtehadi, M. R. Glycan-mediated functional assembly of IL-1RI: structural insights into completion of the current description for immune response. *Journal of Biomolecular Structure and Dynamics* **40**, 2575-2585 (2022).
220 Bering, E., Torstensen, J. Ø., Lervik, A. & de Wijn, A. S. Computational study of the dissolution of cellulose into single chains: the role of the solvent and agitation. *Cellulose* **29**, 1365-1380 (2022).
221 Haji, S., Ito, T., Guenther, C., Nakano, M., Shimizu, T., Mori, D., Chiba, Y., Tanaka, M., Mishra, S. K., Willment, J. A., Brown, G. D., Nagae, M. & Yamasaki, S. Human Dectin-1 is O-glycosylated and serves as a ligand for C-type lectin receptor CLEC-2. *eLife* **11**, e83037, https://doi.org/10.7554/eLife.83037 (2022).
222 Cheatham; III; G.A. Cisneros; V.W.D. Cruzeiro; T.A. Darden; N. Forouzesh; G. Giambaşu; T. Giese; M.K. Gilson; H. Gohlke; A.W. Goetz; J. Harris; S. Izadi; S.A. Izmailov; K. Kasavajhala; M.C. Kaymak; E. King; A. Kovalenko; T. Kurtzman; T.S. Lee; P. Li; C. Lin; J. Liu; T. Luchko; R. Luo; M. Machado; V. Man; M. Manathunga; K.M. Merz; Y. Miao; O. Mikhailovskii; G. Monard; H. Nguyen; K.A. O'Hearn; A. Onufriev; F. Pan; S. Pantano; R. Qi; A. Rahnamoun; D.R. Roe; A. Roitberg; C. Sagui; S. Schott-Verdugo; A. Shajan; J. Shen; C.L. Simmerling; N.R. Skrynnikov; J. Smith; J. Swails; R.C. Walker; J. Wang; J. Wang; H. Wei; X. Wu; Y. Wu; Y. Xiong; Y. Xue; D.M. York; S. Zhao; Q. Zhu; P.A. Kollman, D. A. C. H. M. A. K. B. I. Y. B.-S. J. T. B. S. R. B. D. S. C. T. E. Amber 2023. *University of California, San Francisco* (2023).
223 Abanades, B., Wong, W. K., Boyles, F., Georges, G., Bujotzek, A. & Deane, C. M. ImmuneBuilder: Deep-Learning models for predicting the structures of immune proteins. *Commun Biol* **6**, 575, https://doi.org/10.1038/s42003-023-04927-7 (2023).
224 Hayes, A. J. & Melrose, J. HS, an ancient molecular recognition and information storage glycosaminoglycan, equips hs-proteoglycans with diverse matrix and cell-interactive properties operative in tissue development and tissue function in health and disease. *International Journal of Molecular Sciences* (2023).

225 Liu, X., Yang, Z., Liu, C., Xu, B., Wang, X., Li, Y., Xia, J., Li, D., Zhang, C. & Sun, H. Identification of a type II LacNAc specific binding lectin CMRBL from Cordyceps militaris. *International Journal of Biological Macromolecules* **230**, 123207 (2023).

226 Gruber, C. E. M., Tucci, F. G., Rueca, M., Mazzotta, V., Gramigna, G., Vergori, A., Fabeni, L., Berno, G., Giombini, E. & Butera, O. Treatment-Emergent Cilgavimab Resistance Was Uncommon in Vaccinated Omicron BA. 4/5 Outpatients. *Biomolecules* **13**, 1538 (2023).

227 Jayaprakash, N. G., Sarkar, D. K. & Surolia, A. Atomic visualization of flipped-back conformations of high mannose glycans interacting with cargo lectins: An MD simulation perspective. *Proteins: Structure, Function, and Bioinformatics* (2023).

228 Maity, S. & Acharya, A. Many roles of carbohydrates: A computational spotlight on the coronavirus S protein binding. *ACS Applied Bio Materials* **7**, 646-656 (2023).

229 Carvajal-Barriga, E. J. & Fields, R. D. Sulfated polysaccharides as multi target molecules to fight COVID 19 and comorbidities. *Heliyon* **9** (2023).

230 Roy, R., Poddar, S., Sk, M. F. & Kar, P. Conformational preferences of triantennary and tetraantennary hybrid N-glycans in aqueous solution: Insights from 20 μs long atomistic molecular dynamic simulations. *Journal of Biomolecular Structure and Dynamics* **41**, 3305-3320 (2023).

231 Szklany, K., Mocellin, O., Knippels, L. M., Garssen, J., Kraneveld, A. D. & Mukherjee, R. *Modulation of gut-immune-brain axis by non-digestible oligosaccharides and omega-3 fatty acids in health and allergic disease Synergy or rivalry?* PhD thesis, (2023).

232 Deng, B., Yue, Y., Yang, J., Yang, M., Xing, Q., Peng, H., Wang, F., Li, M., Ma, L. & Zhai, C. Improving the activity and thermostability of PETase from Ideonella sakaiensis through modulating its post-translational glycan modification. *Communications Biology* **6**, 39 (2023).

233 Matamoros-Recio, A., Merino, J., Gallego-Jiménez, A., Conde-Alvarez, R., Fresno, M. & Martín-Santamaría, S. Immune evasion through Toll-like receptor 4: The role of the core oligosaccharides from α2-Proteobacteria atypical lipopolysaccharides. *Carbohydrate Polymers* **318**, 121094 (2023).

234 Bagdonas, H. *Towards prediction of N-glycan compositions from atomic structural data*, University of York, (2023).

235 Flajnik, M. F., Stanfield, R., Pokidysheva, E. N., Boudko, S. P., Wilson, I. & Ohta, Y. An ancient MHC-linked gene encodes a nonrearranging shark antibody, UrIg, convergent with IgG. *The Journal of Immunology* **211**, 1042-1051 (2023).

236 Tatsuoka, H. & Yamaguchi, T. NMR analyses of carbohydrate–water and water–water interactions in water/DMSO mixed solvents, highlighting various hydration behaviors of monosaccharides glucose, galactose and mannose. *BCSJ* **96**, 168-174 (2023).

237 Crawford, C. J., Guazzelli, L., McConnell, S. A., McCabe, O., d'Errico, C., Greengo, S. D., Wear, M. P., Jedlicka, A. E., Casadevall, A. & Oscarson, S. Synthetic glycans reveal determinants of antibody functional efficacy against a fungal pathogen. *ACS infectious diseases* **10**, 475-488 (2023).

238 Leffler, H. Structure and biochemical analysis of galectin interactions with glycoproteins. *Glycoforum* **26**, A17 (2023).

239 Roshini, J., Patro, L. P. P., Sundaresan, S. & Rathinavelan, T. Structural diversity among Acinetobacter baumannii K-antigens and its implication in the in silico serotyping. *Frontiers in Microbiology* **14**, 1191542 (2023).

240 Lossio, C. F., Osterne, V. J., Pinto-Junior, V. R., Chen, S., Oliveira, M. V., Verduijn, J., Verbeke, I., Serna, S., Reichardt, N. C. & Skirtach, A. Structural Analysis and Characterization of an Antiproliferative Lectin from Canavalia villosa Seeds. *International Journal of Molecular Sciences* **24**, 15966 (2023).

241 Samanta, P., Mishra, S. K., Pomin, V. H. & Doerksen, R. J. Docking and Molecular Dynamics Simulations Clarify Binding Sites for Interactions of Novel Marine Sulfated Glycans with SARS-CoV-2 Spike Glycoprotein. *Molecules* **28**, 6413 (2023).
242 Gamarra, M. D., Dieterle, M. E., Capurro, J. I. B., Radusky, L., Piuri, M. & Modenutti, C. P. Molecular Dynamics and Water site bias docking method allows the identification of key amino acids in the Carbohydrate Recognition Domain of a viral protein. *bioRxiv*, 2023.2006. 2001.543333 (2023).
243 Le, H. T., Liu, M. & Grimes, C. L. Application of bioanalytical and computational methods in decoding the roles of glycans in host-pathogen interactions. *Current opinion in chemical biology* **74**, 102301 (2023).
244 Cogez, V., Vicogne, D., Schulz, C., Portier, L., Venturi, G., De Ruyck, J., Decloquement, M., Lensink, M. F., Brysbaert, G. & Dall'Olio, F. N-Glycan on the Non-Consensus NXC Glycosylation Site Impacts Activity, Stability, and Localization of the Sda Synthase B4GALNT2. *International Journal of Molecular Sciences* **24**, 4139 (2023).
245 Trastoy, B., Du, J. J., Cifuente, J. O., Rudolph, L., García-Alija, M., Klontz, E. H., Deredge, D., Sultana, N., Huynh, C. G. & Flowers, M. W. Mechanism of antibody-specific deglycosylation and immune evasion by Streptococcal IgG-specific endoglycosidases. *Nature Communications* **14**, 1705 (2023).
246 Svilenov, H. L., Delhommel, F., Siebenmorgen, T., Rührnößl, F., Popowicz, G. M., Reiter, A., Sattler, M., Brockmeyer, C. & Buchner, J. Extrinsic stabilization of antiviral ACE2-Fc fusion proteins targeting SARS-CoV-2. *Communications Biology* **6**, 386 (2023).
247 Rathore, A. S., Guttman, A., Shrivastava, A. & Joshi, S. Recent progress in high-throughput and automated characterization of N-glycans in monoclonal antibodies. *TrAC Trends in Analytical Chemistry*, 117397 (2023).
248 Guo, Z., Wang, L., Rao, D., Liu, W., Chen, S., Lu, M., Su, L., Chen, S. & Wu, J. Mechanistic Insights into How the Protonation State of D234 Dictates the Reactivity in Streptomyces coelicolor β-N-Acetylhexosaminidase. *The Journal of Physical Chemistry B* **127**, 4820-4828 (2023).
249 Nazipova, A., Makshakova, O. & Kozlova, L. The In Silico Characterization of Monocotyledonous α-L-Arabinofuranosidases on the Example of Maize. *Life* **13**, 266 (2023).
250 Rollins, Z., Harris, B., George, S. & Faller, R. A molecular dynamics investigation of N-glycosylation effects on T-cell receptor kinetics. *Journal of Biomolecular Structure and Dynamics* **41**, 5614-5623 (2023).
251 Muñoz-Basagoiti, J., Monteiro, F. L. L., Krumpe, L. R., Armario-Najera, V., Shenoy, S. R., Perez-Zsolt, D., Westgarth, H. J., Villorbina, G., Bomfim, L. M. & Raïch-Regué, D. Cyanovirin-N binds to select SARS-CoV-2 spike oligosaccharides outside of the receptor binding domain and blocks infection by SARS-CoV-2. *Proceedings of the National Academy of Sciences* **120**, e2214561120 (2023).
252 Di Lorenzo, F., Nicolardi, S., Marchetti, R., Vanacore, A., Gallucci, N., Duda, K., Nieto Fabregat, F., Nguyen, H. N. A., Gully, D. & Saenz, J. Expanding knowledge of methylotrophic capacity: structure and properties of the rough-type lipopolysaccharide from Methylobacterium extorquens and its role on membrane resistance to methanol. *JACS Au* **3**, 929-942 (2023).
253 Eichfeld, R., Mahdi, L. K., De Quattro, C., Armbruster, L., Endeshaw, A. B., Miyauchi, S., Hellmann, M. J., Cord-Landwehr, S., Grigoriev, I. & Peterson, D. Time-resolved transcriptomics reveal a mechanism of host niche defense: beneficial root endophytes deploy a host-protective antimicrobial GH18-CBM5 chitinase. *bioRxiv*, 2023.2012. 2029.572992 (2023).

254 Gass, D. T., Cordes, M. S., Alberti, S. N., Kim, H. J. & Gallagher, E. S. Evidence of H/D Exchange within Metal-Adducted Carbohydrates after Ion/Ion-Dissociation Reactions. *Journal of the American Chemical Society* **145**, 23972-23985 (2023).
255 Costa, G. J., Egbemhenghe, A. & Liang, R. Computational Characterization of the Reactivity of Compound I in Unspecific Peroxygenases. *The Journal of Physical Chemistry B* **127**, 10987-10999 (2023).
256 Wang, Z., Teixeira, S. C., Strother, C., Bowen, A., Casadevall, A. & Cordero, R. J. Neutron Scattering Analysis of Cryptococcus neoformans Polysaccharide Reveals Solution Rigidity and Repeating Fractal-like Structural Patterns. *Biomacromolecules* **25**, 690-699 (2023).
257 Maurya, A. K., Sharma, P., Samanta, P., Shami, A. A., Misra, S. K., Zhang, F., Thara, R., Kumar, D., Shi, D. & Linhardt, R. J. Structure, anti-SARS-CoV-2, and anticoagulant effects of two sulfated galactans from the red alga Botryocladia occidentalis. *International journal of biological macromolecules* **238**, 124168 (2023).
258 Costa, G. J. & Liang, R. Understanding the multifaceted mechanism of Compound I formation in unspecific peroxygenases through multiscale simulations. *The Journal of Physical Chemistry B* **127**, 8809-8824 (2023).
259 Samaniego, L. V. B., Higasi, P. M. R., de Mello Capetti, C. C., Cortez, A. A., Pratavieira, S., Pellegrini, V. d. O. A., Dabul, A. N. G., Segato, F. & Polikarpov, I. Staphylococcus aureus microbial biofilms degradation using cellobiose dehydrogenase from Thermothelomyces thermophilus M77. *International Journal of Biological Macromolecules* **247**, 125822 (2023).
260 Stratilová, B., Šesták, S., Stratilová, E., Vadinová, K., Kozmon, S. & Hrmova, M. Engineering of substrate specificity in a plant cell-wall modifying enzyme through alterations of carboxyl-terminal amino acid residues. *The Plant Journal* **116**, 1529-1544 (2023).
261 Dhurua, S. & Jana, M. Sulfation Effects of Chondroitin Sulfate to Bind a Chemokine in Aqueous Medium: Conformational Heterogeneity and Dynamics from Molecular Simulation. *Journal of Chemical Information and Modeling* **63**, 5660-5675 (2023).
262 Vasudevan, N. K., Li, D. & Xi, L. Potential of Mean Force of Short-Chain Surface Adsorption using Non-Uniform Sampling Windows for Optimal Computational Efficiency. *Macromol. Theory Simul.* (2023).
263 de Mello Capetti, C. C., Pellegrini, V. O. A., Santo, M. C. E., Cortez, A. A., Falvo, M., da Silva Curvelo, A. A., Campos, E., Filgueiras, J. G., Guimaraes, F. E. G. & de Azevedo, E. R. Enzymatic production of xylooligosaccharides from corn cobs: Assessment of two different pretreatment strategies. *Carbohydrate Polymers* **299**, 120174 (2023).
264 Stelfox, A. J., Oguntuyo, K. Y., Rissanen, I., Harlos, K., Rambo, R., Lee, B. & Bowden, T. A. Crystal structure and solution state of the C-terminal head region of the narmovirus receptor binding protein. *Mbio* **14**, e01391-01323 (2023).
265 Dena-Beltrán, J. L., Nava-Domínguez, P., Palmerín-Carreño, D., Martínez-Alarcón, D., Moreno-Celis, U., Valle-Pacheco, M., Castro-Guillén, J. L., Blanco-Labra, A. & García-Gasca, T. EGFR and p38MAPK Contribute to the Apoptotic Effect of the Recombinant Lectin from Tepary Bean (Phaseolus acutifolius) in Colon Cancer Cells. *Pharmaceuticals* **16**, 290 (2023).
266 Luo, R., Jiang, Y., Von Lau, E. & Wu, G. Microscopic influence mechanisms of polysaccharide on the adsorption and corrosion inhibition performance of imidazoline on metal surface. *Applied Surface Science* **613**, 155798 (2023).
267 Jiang, L. & Zheng, K. Xanthoceras sorbifolium Bunge leaf extract activated chia seeds mucilage/chitosan composite film: Structure, performance, bioactivity, and molecular dynamics perspectives. *Food Hydrocolloids* **144**, 109050 (2023).

268 Katara, P., Tyagi, S. & Gupta, M. K. in *Genomics of Plant–Pathogen Interaction and the Stress Response* 1-16 (CRC Press, 2023).
269 Yunpeng, L., Yang, C., Zhao, T., Liu, Y., Liu, Y. & Jiaxue, X. Effect of Temperature on Moisture Adsorption and Desorption in Cellulose Insulation. *Authorea Preprints* (2023).
270 Brunetti, N. S., Davanzo, G. G., de Moraes, D., Ferrari, A. J., Souza, G. F., Muraro, S. P., Knittel, T. L., Boldrini, V. O., Monteiro, L. B. & Virgílio-da-Silva, J. V. SARS-CoV-2 uses CD4 to infect T helper lymphocytes. *Elife* **12**, e84790 (2023).
271 Mani, A. & Kushwaha, S. (CRC Press, 2023).
272 Kim, S. B., Farrag, M., Mishra, S. K., Misra, S. K., Sharp, J. S., Doerksen, R. J. & Pomin, V. H. Selective 2-desulfation of tetrasaccharide-repeating sulfated fucans during oligosaccharide production by mild acid hydrolysis. *Carbohydrate polymers* **301**, 120316 (2023).
273 Capetti, C. C. d. M. *Aplicação de arabinofuranosidase em sinergismo com xilanases para produção de oligossacarídeos a partir de bagaço de cana-de-açúcar* PhD thesis, Universidade de São Paulo, (2023).
274 Abbas, M., Maalej, M., Nieto-Fabregat, F., Thépaut, M., Kleman, J.-P., Ayala, I., Molinaro, A., Simorre, J.-P., Marchetti, R. & Fieschi, F. The unique three-dimensional arrangement of Macrophage Galactose Lectin enables E. coli LipoPolySaccharides recognition through two distinct interfaces. *bioRxiv*, 2023.2003. 2002.530591 (2023).
275 Acosta-Gutiérrez, S., Buckley, J. & Battaglia, G. The Role of Host Cell Glycans on Virus Infectivity: The SARS-CoV-2 Case. *Advanced Science* **10**, 2201853 (2023).
276 Álvarez-Sánchez, M. E., Ruiz-May, E., Aguilar-Tipacamú, G., Elizalde-Contreras, J. M., Bojórquez-Velázquez, E., Zamora-Briseño, J. A., Limón, A. M. V. & Vázquez-Carrillo, L. I. The ovaries of Ivermectin-resistant Rhipicephalus microplus strains display proteomic adaptations involving the induction of xenobiotic detoxification and structural remodeling mechanisms. *Journal of Proteomics* **280**, 104892 (2023).
277 Anso, I., Naegeli, A., Cifuente, J. O., Orrantia, A., Andersson, E., Zenarruzabeitia, O., Moraleda-Montoya, A., García-Alija, M., Corzana, F. & Del Orbe, R. A. Turning universal O into rare Bombay type blood. *Nature Communications* **14**, 1765 (2023).
278 Behren, S., Yu, J., Pett, C., Schorlemer, M., Heine, V., Fischöder, T., Elling, L. & Westerlind, U. Fucose binding motifs on mucin core glycopeptides impact bacterial lectin recognition. *Angewandte Chemie* **135**, e202302437 (2023).
279 Xiao, Y. & Woods, R. J. Protein-Ligand CH-π Interactions: Structural Informatics, Energy Function Development, and Docking Implementation. *J Chem Theory Comput* **19**, 5503-5515, https://doi.org/10.1021/acs.jctc.3c00300 (2023).
280 Meredith, R., Zhu, Y., Yoon, M. K., Tetrault, T., Lin, J., Zhang, W., McGurn, M., Cook, E., Popp, R. & Shit, P. Methyl α-D-galactopyranosyl-(1→ 3)-β-D-galactopyranoside and methyl β-D-galactopyranosyl-(1→ 3)-β-D-galactopyranoside: Glycosidic linkage conformation determined from MA'AT analysis. *Magnetic Resonance in Chemistry* **62**, 544-555 (2024).
281 Zhao, K., Zhang, S., Piao, C., Xu, F., Zhang, Y., Wang, X., Zhang, J., Zhao, C., You, S. G. & Zhang, Y. Investigation of the formation mechanism of the pepper starch-piperine complex. *Int J Biol Macromol* **268**, 131777 (2024).
282 Xu, Z., Zhang, H., Tian, J., Ku, X., Wei, R., Hou, J., Zhang, C., Yang, F., Zou, X. & Li, Y. O-glycosylation of SARS-CoV-2 spike protein by host O-glycosyltransferase strengthens its trimeric structure: O-glycosylation strengthens the trimer of the SARS-CoV-2 S protein. *ABBS* **56**, 1118 (2024).
283 Zhang, W., Meredith, R. J., Yoon, M.-K., Carmichael, I. & Serianni, A. S. Context Effects on Human Milk Oligosaccharide Linkage Conformation and Dynamics Revealed by MA'AT Analysis. *Biochemistry* (2024).

284 Meredith, R. J., Yoon, M.-K., Carmichael, I. & Serianni, A. S. MA'AT Analysis: Unbiased Multi-State Conformational Modeling of Exocyclic Hydroxymethyl Group Conformation in Methyl Aldohexopyranosides. *The Journal of Physical Chemistry B* **128**, 2360-2370 (2024).
285 Nieto-Fabregat, F., Marseglia, A., Thépaut, M., Kleman, J.-P., Abbas, M., Le Roy, A., Ebel, C., Maalej, M., Simorre, J.-P. & Laguri, C. Molecular recognition of Escherichia coli R1-type core lipooligosaccharide by DC-SIGN. *Iscience* **27** (2024).
286 Yoon, M.-K., Shit, P., Zhang, W., Meredith, R. J., Kang, H., Carmichael, I. & Serianni, A. S. 4h J CHOCH Spin Coupling in a Lewisx Trisaccharide as Evidence of Inter-Residue C–H··· O Hydrogen Bonding in Aqueous Solution. *Journal of the American Chemical Society* (2024).
287 Nieto-Fabregat, F., Zhu, Q., Vivès, C., Zhang, Y., Marseglia, A., Chiodo, F., Thépaut, M., Rai, D., Kulkarni, S. S. & Di Lorenzo, F. Atomic-Level Dissection of DC-SIGN Recognition of Bacteroides vulgatus LPS Epitopes. *JACS Au* **4**, 697-712 (2024).
288 Wang, L., Wang, Z., Zhang, H., Jin, Q., Fan, S., Liu, Y., Huang, X., Guo, J., Cai, C. & Zhang, J.-r. A novel esterase regulates Klebsiella pneumoniae hypermucoviscosity and virulence. *PLoS pathogens* **20**, e1012675 (2024).
289 Hao, Z., Li, Z., Zhou, Q., Ma, Z., Lv, J., Wang, Y., Hu, A., Cheng, J., Yu, Z. & Xie, Z. Investigation of the effect of ultrasonication on starch-fatty acid complexes and the stabilization mechanism. *Food Research International* **191**, 114711 (2024).
290 Wang, T. H., Shao, H. P., Zhao, B. Q. & Zhai, H. L. Molecular Insights into the Variability in Infection and Immune Evasion Capabilities of SARS-CoV-2 Variants: A Sequence and Structural Investigation of the RBD Domain. *Journal of Chemical Information and Modeling* **64**, 3503-3523 (2024).
291 Versluys, M., Porras-Domínguez, J. R., Voet, A., Struyf, T. & Van den Ende, W. Insights in inulin binding and inulin oligosaccharide formation by novel multi domain endo-inulinases from Botrytis cinerea. *Carbohydrate Polymers* **328**, 121690 (2024).
292 Lin, Z., Liu, Y., Gong, X., Nie, F., Xu, J. & Guo, Y. Construction of quercetin-fucoidan nanoparticles and their application in cancer chemo-immunotherapy treatment. *International Journal of Biological Macromolecules* **256**, 128057 (2024).
293 Nieto-Fabregat, F., Lenza, M. P., Marseglia, A., Di Carluccio, C., Molinaro, A., Silipo, A. & Marchetti, R. Computational toolbox for the analysis of protein–glycan interactions. *Beilstein Journal of Organic Chemistry* **20**, 2084-2107 (2024).
294 Ranaudo, A., Giulini, M., Pelissou Ayuso, A. & Bonvin, A. M. Modeling Protein–Glycan Interactions with HADDOCK. *Journal of Chemical Information and Modeling* (2024).
295 Widmalm, G. r. Glycan Shape, Motions, and Interactions Explored by NMR Spectroscopy. *JACS Au* **4**, 20-39 (2024).
296 Wei, X., Xie, H., Hu, Z., Zeng, X., Dong, H., Liu, X. & Bai, W. Multiscale structure changes and mechanism of polyphenol-amylose complexes modulated by polyphenolic structures. *International Journal of Biological Macromolecules* **262**, 130086 (2024).
297 Fogarty, C. A., Ives, C. M., Singh, O. & Fadda, E. in *New Developments in Mass Spectrometry* Vol. 15 (ed Weston B Struwe) 381 (Royal Society of Chemistry, 2024).
298 Könighofer, E., Mirgorodskaya, E., Nyström, K., Stiasny, K., Kärmander, A., Bergström, T. & Nordén, R. Identification of Three Novel O-Linked Glycans in the Envelope Protein of Tick-Borne Encephalitis Virus. (2024).
299 Ives, C. M., Singh, O., D'Andrea, S., Fogarty, C. A., Harbison, A. M., Satheesan, A., Tropea, B. & Fadda, E. Restoring protein glycosylation with GlycoShape. *Nature Methods*, 1-11 (2024).
300 Unione, L., Ammerlaan, A. N., Bosman, G. P., Uslu, E., Liang, R., Broszeit, F., van der Woude, R., Liu, Y., Ma, S. & Liu, L. Probing altered receptor specificities of antigenically

drifting human H3N2 viruses by chemoenzymatic synthesis, NMR, and modeling. *Nature Communications* **15**, 2979 (2024).
301 Ulloa-Aguirre, A., Llamosas, R. & Dias, J. A. Follicle-stimulating hormone sweetness: How carbohydrate structures impact on the biological function of the hormone. *Archives of Medical Research* **55**, 103091 (2024).
302 Yang, T., Zhang, Y., Guo, L., Li, D., Liu, A., Bilal, M., Xie, C., Yang, R., Gu, Z. & Jiang, D. Antifreeze polysaccharides from wheat bran: The structural characterization and antifreeze mechanism. *Biol. Macromol.* (2024).
303 Tiemblo-Martín, M., Pistorio, V., Saake, P., Mahdi, L., Campanero-Rhodes, M. A., Di Girolamo, R., Di Carluccio, C., Marchetti, R., Molinaro, A. & Solís, D. Structure and properties of the exopolysaccharide isolated from Flavobacterium sp. Root935. *Carbohydrate polymers* **343**, 122433 (2024).
304 Osada, N., Mishra, S. K., Nakano, M., Tokoro, Y., Nagae, M., Doerksen, R. J. & Kizuka, Y. Self-regulation of MGAT4A and MGAT4B activity toward glycoproteins through interaction of lectin domain with their own N-glycans. *iScience* **27** (2024).
305 Li, L., Wu, D., Gui, H., Gao, T., Jing, H., Duan, N., Wang, S., Liu, L., Shen, Y. & Xu, Z. Interaction Mechanism between Cyanidin-3-O-Glucoside and Different Types of Corn Starch and Their Effect on the Multi-Scale Structure of Starch. *Available at SSRN 4861700* (2024).
306 Pinto-Junior, V. R., Seeger, R. L., Souza-Filho, C. H. D., França, A. P., Sartori, N., Oliveira, M. V., Osterne, V. J. S., Nascimento, K. S., Leal, R. B. & Cavada, B. S. Xc-System as a Possible Target for ConBr Lectin Interaction in Glioma Cells. *Neuroglia* **5**, 202-222 (2024).
307 Cinar, M. S., Niyas, A. & Avci, F. Y. Serine-rich repeat proteins: well-known yet little-understood bacterial adhesins. *Journal of Bacteriology* **206**, e00241-00223 (2024).
308 Riopedre-Fernandez, M., Kostal, V., Martinek, T., Martinez-Seara, H. & Biriukov, D. Developing and Benchmarking Sulfate and Sulfamate Force Field Parameters via Ab Initio Molecular Dynamics Simulations To Accurately Model Glycosaminoglycan Electrostatic Interactions. *Journal of Chemical Information and Modeling* **64**, 7122-7134 (2024).
309 Taniguchi, K., Karita, S. & Umekawa, M. Synergistic xylan decomposition by a reducing-end xylose-releasing exo-oligoxylanase with other xylanolytic enzymes derived from Paenibacillus xylaniclasticus strain TW1. *Bioscience, Biotechnology, and Biochemistry*, zbae130 (2024).
310 Vacilotto, M. M., de Araujo Montalvão, L., Pellegrini, V. d. O. A., Liberato, M. V., de Araujo, E. A. & Polikarpov, I. Two-domain GH30 xylanase from human gut microbiota as a tool for enzymatic production of xylooligosaccharides: Crystallographic structure and a synergy with GH11 xylosidase. *Carbohydrate Polymers* **337**, 122141 (2024).
311 Urban, J., Jin, C., Thomsson, K. A., Karlsson, N. G., Ives, C. M., Fadda, E. & Bojar, D. Predicting glycan structure from tandem mass spectrometry via deep learning. *Nature Methods* **21**, 1206-1215 (2024).
312 Riopedre-Fernandez, M., Kostal, V., Martinek, T., Martinez-Seara, H. & Biriukov, D. Developing and Benchmarking Sulfate and Sulfamate Force Field Parameters for Glycosaminoglycans via Ab Initio Molecular Dynamics Simulations. *bioRxiv*, 2024.2005. 2031.596767 (2024).
313 Schachner, L. F., Mullen, C., Phung, W., Hinkle, J. D., Beardsley, M. I., Bentley, T., Day, P., Tsai, C., Sukumaran, S. & Baginski, T. Exposing the molecular heterogeneity of glycosylated biotherapeutics. *Nature Communications* **15**, 3259 (2024).
314 Gazi, I., Reiding, K. R., Groeneveld, A., Bastiaans, J., Huppertz, T. & Heck, A. J. LacdiNAc to LacNAc: remodelling of bovine α-lactalbumin N-glycosylation during the transition from colostrum to mature milk. *Glycobiology* **34**, cwae062 (2024).

315 Gamarra, M. D., Dieterle, M. E., Ortigosa, J., Lannot, J. O., Blanco Capurro, J. I., Di Paola, M., Radusky, L., Duette, G., Piuri, M. & Modenutti, C. P. Unveiling crucial amino acids in the carbohydrate recognition domain of a viral protein through a structural bioinformatic approach. *Glycobiology* **34**, cwae068 (2024).
316 Briganti, L., Manzine, L. R., de Mello Capetti, C. C., de Araújo, E. A., Pellegrini, V. d. O. A., Guimaraes, F. E. G., de Oliveira Neto, M. & Polikarpov, I. Unravelling biochemical and structural features of Bacillus licheniformis GH5 mannanase using site-directed mutagenesis and high-resolution protein crystallography studies. *International Journal of Biological Macromolecules*, 133182 (2024).
317 Dey, R. & Taraphder, S. Molecular Modeling of Glycosylated Catalytic Domain of Human Carbonic Anhydrase IX. *The Journal of Physical Chemistry B* (2024).
318 Gimeno, A., Ehlers, A. M., Delgado, S., Langenbach, J.-W. H., van den Bos, L. J., Kruijtzer, J. A., Guigas, B. G. & Boons, G.-J. A Chemoenzymatic Strategy for Site-Specific Glyco-Tagging of Native Proteins for the Development of Biologicals. *bioRxiv*, 2024.2008. 2013.607754 (2024).
319 De Souza, T. P., Cantão, L. X. S., Rodrigues, M. Q. R. B., Gonçalves, D. B., Nagem, R. A. P., Rocha, R. E. O., Godoi, R. R., Lima, W. J. N., Galdino, A. S. & Minardi, R. C. d. M. Glycosylation and charge distribution orchestrates the conformational ensembles of a biotechnologically promissory phytase in different pHs–a computational study. *Journal of Biomolecular Structure and Dynamics* **42**, 5030-5041 (2024).
320 Hafeez, S. & Zaidi, N. U. S. S. Prevention of Blood Incompatibility Related Hemagglutination: Blocking of Antigen A on Red Blood Cells Using In Silico Designed Recombinant Anti-A scFv. *Antibodies* **13**, 64 (2024).
321 Quintana, J. I., Delgado, S., Rábano, M., Azkargorta, M., Florencio-Zabaleta, M., Unione, L., Vivanco, M. d., Elortza, F., Jiménez-Barbero, J. & Ardá, A. The impact of glycosylation on the structure, function, and interactions of CD14. *Glycobiology* **34**, cwae002 (2024).
322 Dignon, G. L. & Dill, K. A. Computational Procedure for Predicting Excipient Effects on Protein–Protein Affinities. *Journal of Chemical Theory and Computation* **20**, 1479-1488 (2024).
323 Lundstrøm, J., Gillon, E., Chazalet, V., Kerekes, N., Di Maio, A., Feizi, T., Liu, Y., Varrot, A. & Bojar, D. Elucidating the glycan-binding specificity and structure of Cucumis melo agglutinin, a new R-type lectin. *Beilstein Journal of Organic Chemistry* **20**, 306-320 (2024).
324 Prestegard, J. H. A consensus structural motif for the capsular polysaccharide of Cryptococcus Neoformans by NMR/MD. *Proceedings of the National Academy of Sciences* **121**, e2322413121 (2024).
325 Tan, Z., Ning, L., Cao, L., Zhou, Y., Li, J., Yang, Y., Lin, S., Ren, X., Xue, X. & Kang, H. Bisecting GlcNAc modification reverses the chemoresistance via attenuating the function of P-gp. *Theranostics* **14**, 5184 (2024).
326 Dhurua, S., Maity, S., Maity, B. & Jana, M. Comparative Bindings of Glycosaminoglycans with CXCL8 Monomer and Dimer: Insights from Conformational Dynamics and Kinetics of Hydrogen Bonds. *The Journal of Physical Chemistry B* (2024).
327 Wang, D., Zhang, Z., Baudys, J., Haynes, C., Osman, S. H., Zhou, B., Barr, J. R. & Gumbart, J. C. Enhanced surface accessibility of SARS-CoV-2 Omicron spike protein due to an altered glycosylation profile. *ACS Infectious Diseases* (2024).
328 Zong, Y., Lei, Z., Yu, S.-B., Zhang, L.-Y., Wu, Y., Feng, K., Qi, Q.-Y., Liu, Y., Zhu, Y. & Guo, P. Caltrop-like Small-Molecule Antidotes That Neutralize Unfractionated Heparin and Low-Molecular-Weight Heparin In Vivo. *J Med Chem* **67**, 3860-3873 (2024).
329 Enmozhi, S. K., Raja, K., Kaleeswaran, M., Vivek, R., Samuel, R. S., Sebastine, I., Murugan, R., Joseph, J., Chinnasamy, A. & Kaleeswaran, M. Identification of Promising

SARS-CoV-2 Main Protease (Mpro) and Spike Protein Inhibitors From Edible Mushroom: A Computational Approach. *Authorea Preprints* (2024).

330 Dhurua, S. & Jana, M. Conformational preferences of heparan sulfate to recognize the CXCL8 dimer in aqueous medium: degree of sulfation and hydrogen bonds. *Physical Chemistry Chemical Physics* **26**, 21888-21904 (2024).

331 Murphy, P. V., Dhara, A., Fitzgerald, L. S., Hever, E., Konda, S. & Mandal, K. Small lectin ligands as a basis for applications in glycoscience and glycomedicine. *Chemical Society Reviews* (2024).

332 Zariñán, T., Espinal-Enriquez, J., De Anda-Jáuregui, G., Lira-Albarrán, S., Hernández-Montes, G., Gutiérrez-Sagal, R., Rebollar-Vega, R. G., Bousfield, G. R., Butnev, V. Y. & Hernández-Lemus, E. Differential effects of follicle-stimulating hormone glycoforms on the transcriptome profile of cultured rat granulosa cells as disclosed by RNA-seq. *Plos one* **19**, e0293688 (2024).

333 Jager, S., Zeller, M., Pashkova, A., Schulte, D., Damoc, E., Reiding, K., Makarov, A. A. & Heck, A. J. Narrow window data-independent acquisition on the Orbitrap Astral Mass Spectrometer enables fast and deep coverage of the plasma glycoproteome. *bioRxiv*, 2024.2007. 2029.605591 (2024).

334 Osterne, V. J., Pinto-Junior, V. R., Oliveira, M. V., Nascimento, K. S., Van Damme, E. J. & Cavada, B. S. Computational insights into the circular permutation roles on ConA binding and structural stability. *Current Research in Structural Biology* **7**, 100140 (2024).

335 Kil, Y., Pichkur, E. B., Sergeev, V. R., Zabrodskaya, Y., Myasnikov, A., Konevega, A. L., Shtam, T., Samygina, V. R. & Rychkov, G. N. The archaeal highly thermostable GH 35 family β-galactosidase D aβ G al has a unique seven domain protein fold. *The FEBS Journal* (2024).

336 Day, C. J., Favuzza, P., Bielfeld, S., Haselhorst, T., Seefeldt, L., Hauser, J., Shewell, L. K., Flueck, C., Poole, J. & Jen, F. E.-C. The essential malaria protein PfCyRPA targets glycans to invade erythrocytes. *Cell Reports* **43** (2024).

337 Cabeza, O. I., Parra, N., Cerro, R., Mansilla, R., Sanchez, R. Z., Gutierrez-Reinoso, M., Escribano, E. H., Castillo, R., Rodriguez-Alvarez, L. & Tavares, K. Development and characterization of a novel variant of long-acting bovine follicle-stimulating hormone (brscFSH). *Theriogenology* **226**, 76-86 (2024).

338 Nilchan, N., Kraivong, R., Luangaram, P., Phungsom, A., Tantiwatcharakunthon, M., Traewachiwiphak, S., Prommool, T., Punyadee, N., Avirutnan, P. & Duangchinda, T. An Engineered N-Glycosylated Dengue Envelope Protein Domain III Facilitates Epitope-Directed Selection of Potently Neutralizing and Minimally Enhancing Antibodies. *ACS Infectious Diseases* **10**, 2690-2704 (2024).

339 Friganović, T., Borko, V. & Weitner, T. Protein sialylation affects the pH-dependent binding of ferric ion to human serum transferrin. *Dalton Transactions* (2024).

340 de Mello Capetti, C. C., Ontañon, O., Navas, L. E., Campos, E., Simister, R., Dowle, A., Liberato, M. V., Pellegrini, V. d. O. A., Gómez, L. D. & Polikarpov, I. Sugarcane bagasse derived xylooligosaccharides produced by an arabinofuranosidase/xylobiohydrolase from Bifidobacterium longum in synergism with xylanases. *Carbohydrate Polymers* **339**, 122248 (2024).

341 Wang, Z. *Multifaceted Investigation into Cryptococcus neoformans Biomolecules* MS thesis, Johns Hopkins University, (2024).

342 Dolinska, M. B. & Sergeev, Y. V. Molecular Modeling of the Multiple-Substrate Activity of the Human Recombinant Intra-Melanosomal Domain of Tyrosinase and Its OCA1B-Related Mutant Variant P406L. *International Journal of Molecular Sciences* **25**, 3373 (2024).

343 Mehta, D. & Sanhueza, C. A. Interglycosidic C5–C6 rotamer distributions of alkyl O-rutinosides. *Carbohydrate Research* **544**, 109251 (2024).

344 Amos, B. K., Pook, V., Prates, E., Stork, J., Shah, M., Jacobson, D. A. & DeBolt, S. Discovery and Characterization of Fluopipamine, a Putative Cellulose Synthase 1 Antagonist within Arabidopsis. *Journal of Agricultural and Food Chemistry* **72**, 3171-3179 (2024).

345 Bloch, Y., Osterne, V. J., Savvides, S. N. & Van Damme, E. J. The crystal structure of Nictaba reveals its carbohydrate-binding properties and a new lectin dimerization mode. *Glycobiology*, cwae087 (2024).

346 Vasudevan, N. K., Li, D. & Xi, L. Potential of Mean Force of Short-Chain Surface Adsorption using Non-Uniform Sampling Windows for Optimal Computational Efficiency. *Macromolecular Theory and Simulations* **33**, 2300057, https://doi.org/https://doi.org/10.1002/mats.202300057 (2024).

347 Chernykh, A., Abrahams, J. L., Grant, O. C., Kambanis, L., Sumer-Bayraktar, Z., Ugonotti, J., Kawahara, R., Corcilius, L., Payne, R. J., Woods, R. J. & Thaysen-Andersen, M. Position-specific N- and O-glycosylation of the reactive center loop impacts neutrophil elastase-mediated proteolysis of corticosteroid-binding globulin. *J Biol Chem* **300**, 105519, https://doi.org/10.1016/j.jbc.2023.105519 (2024).

348 Zhang, W., Meredith, R. J., Wang, X., Woods, R. J., Carmichael, I. & Serianni, A. S. Does Inter-Residue Hydrogen Bonding in β-(1→4)-Linked Disaccharides Influence Linkage Conformation in Aqueous Solution? *The Journal of Physical Chemistry B* **128**, 2317-2325, https://doi.org/10.1021/acs.jpcb.3c07448 (2024).

349 Cisar, J. O., Wang, X., Woods, R. J., Cain, K. D. & Wiens, G. D. Structural and genetic basis for the binding of a mouse monoclonal antibody to Flavobacterium psychrophilum lipopolysaccharide. *Journal of Fish Diseases* **n/a**, e13958, https://doi.org/https://doi.org/10.1111/jfd.13958 (2024).
